# Long Amplicon Nanopore Sequencing for Dual-Typing *RdRp* and *VP1* Genes of Norovirus Genogroups I and II in Wastewater

**DOI:** 10.1101/2024.03.27.584784

**Authors:** G. Scott, D. Ryder, M. Buckley, R. Hill, S. Treagus, T. Stapleton, D. I. Walker, J. Lowther, F. M. Batista

## Abstract

Noroviruses (NoV) are the leading cause of non-bacterial gastroenteritis across the globe with societal costs of US$60.3 billion per annum. Development of a long amplicon nanopore-based method for dual-typing the *RNA-dependent RNA polymerase* (*RdRp*) and major structural protein (*VP1*) regions from a single RNA fragment could improve existing norovirus typing methods. Its application to wastewater-based epidemiology (WBE) and environmental testing could enable the discovery of novel types and improve tracking throughout the population and into aquaculture and recreational water settings. Here, we develop and optimise such a method for wastewater as the sample matrix.

Reverse transcription (RT), PCR and library pooling were optimised and a consensus-based bioinformatics pipeline was developed. Inhibitor removal and LunaScript® RT gave robust amplification of the ≈1000 bp *RdRP*+*VP1* amplicon. Platinum™ Taq polymerase showed good sensitivity and reduced levels non-specific amplification (NSA) when compared to other polymerases. Optimised PCR annealing temperatures significantly reduced NSA (51.3% and 42.4% for GI and GII), increased yield (86.5% for GII) and increased taxa richness (57.7%) for GII. Analysis of three NoV positive faecal samples showed 100% nucleotide similarity with Sanger sequencing.

Eight GI genotypes, 11 polymerase types (p-types) and 13 combinations were detected in wastewater along with 4 GII genotypes, 4 p-types and 8 combinations; highlighting the diversity of norovirus taxa present in wastewater in England. The most common genotypes detected in clinical samples were all detected in wastewater while we also commonly detected several GI genotypes not reported in the clinical data. Application of this method into a WBE scheme, therefore, may allow for more accurate measurement of norovirus diversity within the population.

## 1.0 Introduction

Noroviruses (NoV) are non-enveloped viruses in the family Caliciviridae with a single stranded, positive sense RNA genome. In human NoV strains the genome is ≈7.6 kb in length and comprised of three open reading frames (ORFs). ORF-1 encodes the non-structural proteins P48, NTPase, P22, VPg, Protease and RNA-dependent RNA polymerase (RdRp) while ORF-2 and -3 encodes the major structural capsid protein VP1 and minor structural protein VP2 (Campillay-Véliz *et al*., 2020). Three of the ten NoV genogroups (GI, GII and GIV) cause gastroenteritis in humans (Baldridge, Turula and Wobus, 2016). Their further classification into genotypes and polymerase-types (p-type) is based on *VP1* and *RdRp* regions, with a dual-typing system (genotype + p-type) proposed by Chhabra *et al*., (2019) naming of both the genotype (e.g. GI.2) and p-type (e.g. GI.P2) in combination (e.g. GI.2[P2]) referred to herein as type.

The leading cause of epidemic and non-bacterial gastroenteritis, NoV was estimated to cost US$4.2 billion in direct health costs and US$60.3 billion in societal costs worldwide in 2016 (Baldridge, Turula and Wobus, 2016; Bartsch *et al*., 2016; Campillay-Véliz *et al*., 2020). Although severe disease and death is rare, mortality rates are higher in children <5 y old, the elderly and immunocompromised (Baldridge, Turula and Wobus, 2016; Bartsch *et al*., 2016). Out of an estimated 699 million illnesses, 219,000 deaths are predicted to occur annually; 70,000 being children <5 y old (Baldridge, Turula and Wobus, 2016; Bartsch *et al*., 2016). The main transmission route is faecal-oral and an acute symptomatic phase of 1 to 4 d occurs with faecal shedding commencing at 0.8 d and may continue for months (Baldridge, Turula and Wobus, 2016; Ge *et al*., 2023). High viral loads (∼2.0 x 10^9^ genome copies/g stool) and particle stability make NoV a suitable candidate for wastewater based epidemiology (WBE) (Hall, 2012; Newman *et al*., 2016; Hassard *et al*., 2017).

WBE became prominent during the SARS-CoV-2 pandemic with at least 72 countries adopting it to quantify viral RNA, track variants of concern and emerging variants (Naughton *et al*., 2021). WBE, however, has been used for decades as part of the poliovirus eradication programme and for monitoring other pathogens such as hepatitis viruses and norovirus (Asghar *et al*., 2014; Polo *et al*., 2020; Treagus *et al*., 2023). It has the potential to be a useful tool with benefits including detection of asymptomatic cases, access to near population-scale epidemiological information without mass testing and fewer anthropogenic biases.

WBE studies have already shown that NoV quantities in wastewater can be predictive (2 to 4 weeks prior) or are coincident with clinical cases (Hellmér *et al*., 2014; Kazama *et al*., 2017; Alex-sanders *et al*., 2023; Boehm *et al*., 2023; Markt *et al*., 2023). Metagenomic and amplicon sequencing approaches have been used to type NoVs. Strubbia *et al*., (2019) detected six types GI.4[P4], GI.2[P2], GI.1[P1], GII.6[P7] GII.17[P17] and GII.4 Sydney [P16] from three composite samples from a city with population of ≈0.3 million using a metagenomic approach paired using MiSeq 2x150 bp (Illumina, USA). Six full-length genomes (>7000nt) were recovered highlighting the benefit of a metagenomic approach which won’t be affected by PCR bias but may suffer from reduced sensitivity.

Numerous studies using amplicon sequencing to genotype and p-type NoV in wastewater have been performed. The majority amplify partial *VP1* or *RdRp* regions in isolation. Kazama *et al*., (2017) and Cao *et al*., (2022) used semi-nested PCR of partial GI and GII *VP1* regions (≈337 bp) using the GS-Junior (Roche, Switzerland) and NovaSeq (Illumina, USA) platforms. In both cases 15 genotypes were detected, with up to 8 genotypes per sample for the former. Fumian *et al*., (2019) amplified the 5’ of the GII *VP1* (373 bp) using MiSeq 2x150 bp (Illumina, USA). Over 1 yr, 13 genotypes were detected from 156 samples with GII.4 being the most prevalent. Mabasa *et al*., (2022) sequenced ≈575bp of the ORF-1 and -2 junction using MiSeq 2x300 bp (Illumina, USA). Over 26 months of bi-weekly sampling, 81% were positive and 13 and 21 types of GI and GII were detected. Recently our group used nanopore sequencing to genotype GII (302 bp amplicon) and found 8 genotypes across 42 samples (Treagus *et al*., 2023). These previous studies highlight the diverse range of NoV types that can be detected in wastewater.

Most previous research, however, has relied on short-read sequencing which often require separate amplification and sequencing of *VP1* and *RdRp.* This makes application of dual-typing post-sequencing extremely difficult due to high levels of recombination in the norovirus genome. The aim of this study, therefore, was to develop a long-amplicon (≈1000bp) nanopore sequencing method optimised for wastewater that allows dual-typing of GI and GII. Development of such an approach could allow identification of emerging variants and novel recombinants from wastewater.

## 2.0 Materials and Methods

A step-by-step protocol for the methods developed in this study is available in the supplementary protocol or on protocols.io at dx.doi.org/10.17504/protocols.io.8epv5xpmjg1b/v1.

### 2.1 Sample Collection and Processing

Samples were collected and processed by the Environment Agency as part of the SARS-CoV-2 wastewater monitoring programme in England (UK) as outlined in Walker, (2024). Briefly, 1 L of untreated sewage was collected and processed (150 mL) using ammonium sulphate precipitation and nucleic acid extraction using a Kingfisher Flex™ (Thermo Scientific™, UK) and NucliSENS® (BioMérieux, France) reagents. Nucleic acids were sent on dry ice to the Centre for Environment, Fisheries and Aquaculture Science (Weymouth, UK).

To assess the performance of inhibitor removal and the wastewater optimised sequencing protocol, 209 nucleic acid extracts from wastewater collected between 22/10/2021 and 25/10/2021 were pooled by geographical region based on the location of the wastewater treatment plants using QGIS 3.16 and the “Regions (December 2021) EN BFC” (Office for National Statistics, 2021; QGIS Development Team, 2021). Regions with limited or excess nucleic acid volume were split, combined or omitted creating 10 samples, Supplementary Table 1. Excess nucleic acids were pooled into three additional independent pools and used to assess reverse transcription, PCR and size selection.

### 2.2 Inhibitor Removal

To assess the impact of inhibitors on reverse transcription and PCR, nucleic acid extracts were cleaned with Mag-Bind® TotalPure NGS beads (Omega Bio-Tek, USA) following Child *et al*., (2023) using 25 µL of nucleic acids. To monitor inhibitor removal efficacy, a GI RT-qPCR and an external control RNA (EC RNA) method was used following ISO 15216-1: 2017 (ISO, 2017). Reactions were spiked with 1 µL of EC RNA (4,000 gc/µL) and run alongside an EC RNA + water control. Reactions were run in duplicate on QuantStudio™ 3 machines. RT-qPCR standard curve slopes were between −3.6 and −3.1 with R^2^≥0.99. Technical repeats were averaged prior to analysis. Inhibition values <0% were assigned a value of 0 and NoV concentration data was square root transformed (West, 2022).

### 2.3 Reverse Transcription Optimisation

Methods using two reverse transcription (RT) kits were tested; SuperScript™ IV Reverse Transcriptase (Invitrogen™, USA) and LunaScript® RT SuperMix (New England Biolabs, USA), referred to as Superscript™ and LunaScript®. For SuperScript™, the manufacturer’s instructions were followed for a 20 µL final volume, 10 µL of nucleic acids and RT at 50°C. LunaScript® followed Child et al., (2023) with a nucleic acid volume of 10 µL. Molecular biology grade water (7.5 µL) was added after RT to equalise sample dilution.

To assess RT performance, semi-nested PCR of the *RdRp+VP1* region was performed. First round and semi-nested products were 1194 and 1110 bp for GI and 1052 and 971 bp for GII, respectively (Table 1). PCR with Platinum™ Taq DNA Polymerase followed the manufacturer’s instructions for a 25 μl reaction with 5 μl of cDNA or first-round PCR products. Cycling conditions were 95°C for 1 min followed by 40 cycles of 95°C for 30 s, 50°C for 30 s and 72°C for 30 s and 72°C for 7 min. Products were visualised by gel electrophoresis with 2% tris-borate EDTA agarose (Sigma Aldrich) with 100 bp ladder (Promega, USA) or with a TapeStation 4150 using D5000 screen tape (Agilent, USA).

**Table 1.**
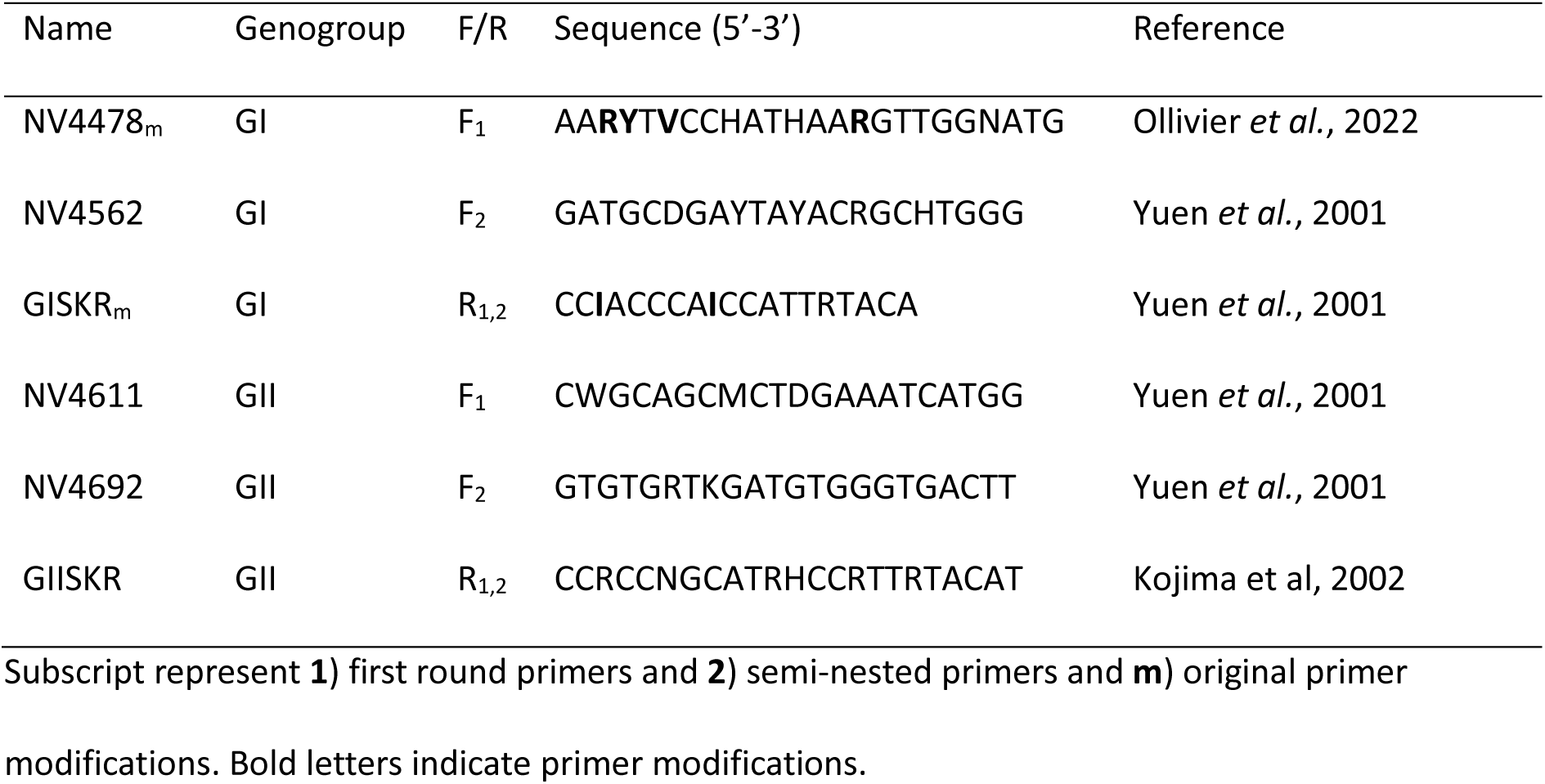
Primers used for amplification of the RdRp+VP1 region.

### 2.4 PCR Optimisation and Size Selection

Two independent pooled cDNA samples processed using LunaScript™ were used to optimise PCR. Six polymerases were assessed; LongAmp® Hot Start (New England Biolabs, USA); Phusion™ Hot Start II DNA Polymerase (Invitrogen™, USA); Platinum™ Taq polymerase (Invitrogen™, USA); Q5U® Hot Start High-Fidelity DNA Polymerase (New England Biolabs, USA); NEB Next® Ultra™ II Q5® mastermix (New England Biolabs, USA); and Platinum™ SuperFi™ DNA Polymerase (Invitrogen™, USA) referred to herein as LongAmp®, Phusion™, Platinum®, Q5U®, Ultra™ II Q5 and SuperFi™. Annealing temperatures (T_a_) were optimised with gradient PCR using the lowest primer melting temperature (T_m_) for each polymerase as the highest T_a_ with two additional temperatures at ≈2.5°C and ≈5.0°C below. Cycling conditions were as recommended by the manufacturer and run for 40 cycles. Reactions were run in simplex at 1 and 10-fold dilutions on a Mastercycler® Nexus Gradient (Eppendorf, Germany). Optimised PCR T_a_ were compared to T_a_=50°C as used by Ollivier et al. (2022). Ampure XP (Beckman Coulter, USA) and Mag-Bind® TotalPure NGS (Omega Bio-Tek, USA) beads were trialled for size selection using both 0.4 to 0.6x ratios based on the manufacturers’ recommendations.

### 2.5 Library Preparation and Sequencing

To simultaneously type GI and GII under a single barcode, an amplicon pooling method was developed to increase the equality of sequencing depth. Nucleic acid extracts (158) from untreated sewage samples collected between 05/08/21 and 11/08/21 and 25/02/22 and 07/03/2022 were processed using the wastewater optimised methods then analysed by TapeStation. The amplicon of interest percentage AOI% was calculated following Supplementary Equation 1. Negative and diluted samples, those with unusually high levels of non-specific amplification (NSA) and outliers (1.5 times the interquartile range) were removed and the average GI:GII AOI% ratio was determined (Supplementary Equation 2). This was used to adjust the moles of the GI and GII PCR products input into library preparation; Supplementary Equations 3 and 4.

PCR products were purified using ExoSAP-IT™ (Applied Biosystems, USA) following the manufacturer’s instructions for 10 µL of product. PCR yield was determined using a Qubit™ Flex (Invitrogen™, USA) and dsDNA High Sensitivity kit with 2 µL of sample. For the wastewater optimised assay GI and GII amplicon pooling, molarities were adjusted for amplicon length and NSA with 85.3 and 114.7 fmol of GI and GII amplicons pooled together. For the unoptimised method, 62.0 and 138.0 fmol of GI and GII were pooled together based on the GI:GII AOI% ratios of the samples under investigation. Library preparation was performed using the Native Barcoding Kit 96 V14 (Oxford Nanopore Technologies, UK) following the manufacturer’s instructions for sequencing of amplicons. Sequencing was performed on a GridION (MinKNOW software release 22.10.7) using R10.4.1 flow cells at 260 bps with super accurate basecalling. Whole process (cDNA synthesis onwards) and PCR-specific no template controls were sequenced alongside the samples.

### 2.6 Bioinformatics

Briefly, reads were split using duplex_tools 0.2.14 and trimmed with cutadapt 3.4 (Martin, 2011; Oxford Nanopore Technologies, 2022). Size filtering and random sampling (>800 bp and ≤90,000 reads per sample) was performed using SeqTK 1.3 (Hang Li, 2018). Minimap 2.24 was used to find overlaps between reads. Alignments were screened using yacrd 1.0.0 to detect chimeras or poorly supported reads where more than 20% of a given read had a depth of less than 10 (Li, 2018; Marijon, Chikhi and Varré, 2020). NGSpeciesID 0.1.3 clustered reads and formed consensus sequences supported by more than 100 reads (Sahlin, Lim and Prost, 2021).

For each sample, consensus sequences were indexed, polished and screened for regions with poor support and reads aligned using kma 1.4.9 (Clausen, Aarestrup and Lund, 2018). All consensus sequences were concatenated, renamed using SeqTK 1.3 and then indexed with Samtools 1.13 (Hang Li, 2018; Danecek *et al*., 2021). SeqKit 2.3.0 identified regions at consensus termini soft masked by kma 1.4.9 due to having poor support, with any such regions being trimmed using Bedtools 2.30.0 (Quinlan and Hall, 2010; Shen *et al*., 2016). Consensuses were clustered at 95% (CD-HIT 4.8.1) and typed using the Centre for Disease Control’s (CDC) Human Calicivirus Typing Tool (Fu *et al*., 2012; Tatusov *et al*., 2021). Full bioinformatic procedures, tools and commands can be found in the Supplementary Table 2.

Untyped consensus sequences were aligned against CDC reference sequences in MEGA 11.0.11 using the ClustalW algorithm with default parameters (Tamura et al., 2021; Centre for Disease Control, 2023b). Indels causing frame-shift mutations were assessed using amino acid alignments and putative viral typing using the CDC’s NoV typing tool followed by errors correction. R 4.1.2 was used to collate the number of reads mapped to each of the consensus sequences (R Core Team, 2021).

Putative PCR chimeras were removed from the dataset if they met any of the following criteria: Failing to align against the CDC’s reference sequence database (Centre for Disease Control, 2023b), identification as a chimera by USEARCH v11 de-novo or a child sequence with a parent breakpoint within the terminal or proximal regions of *RdRp* and *VP1* and child-parent sequence similarities ≥95%. For USEARCH chimera detection, consensus sequences were annotated with read count (R) and screened using USEARCH v11 de-novo chimera detection (Edgar, 2016). Manually screening was performed using NCBI Multiple Sequence Alignment Viewer v1.25.0. For the method comparison study, a read-depth threshold of 0.1% of the median reads per sample was used and reads from consensus sequences of the same viral type were grouped.

Nanopore-generated sequences were assessed against those obtained by Sanger sequencing. Three norovirus positive faecal samples as determined by RT-qPCR following ISO 15216-1:2017 from presumed single-type infections (ISO, 2017), were processed following the wastewater optimised protocol. PCR products were purified using the QIAquick PCR Purification Kit (QIAGEN, Germany) and sequenced with the forward primer using Mix2Seq (Eurofins, Germany). Sequences were trimmed to 10 consecutive Q30 bases prior to alignment. The above bioinformatic approach was used without consensus sequence clustering at 95%. Read quality was estimated by aligning reads against the faecal samples consensus sequences using minimap 2.26 and the map-ont preset. Results were filtered to include only those alignments ≥700 bp. Quality scores were calculated as described in Supplementary Equation 5.

### 2.7 Data Analysis

For all inferential statistics, paired t-tests were performed unless the assumption of normality of the difference between observations was not met where a Wilcoxon signed-rank test was performed. All data analysis and visualisation was performed in R Studio build 353 (2022.12.0) and statistical significance is defined as p<0.05 (Posit team, 2022).

## 3.0 Results

### 3.1 Inhibitor Removal and Reverse Transcription

Inhibitor removal significantly reduced average inhibition from 90.6% to 13.2% and increased NoV quantification 4.8-fold; p<0.001 (Figure 1 A and B). Reverse transcription studies showed no or weak amplification with SuperScript™ while Lunascript® showed good amplification for GI and GII with inhibitor removal (Figure 2 A and B). For Lunascript®, the number of aligned reads ≥900 bp (log_10_) significantly increased with inhibitor removal from 1.99 ± 0.25 (sd) to 5.22 ± 0.24 for GI and 2.31 ± 0.38 to 4.52 ± 0.46 for GII (p<0.001) giving 163-fold and 1731-fold differences.

**Figure 1.**
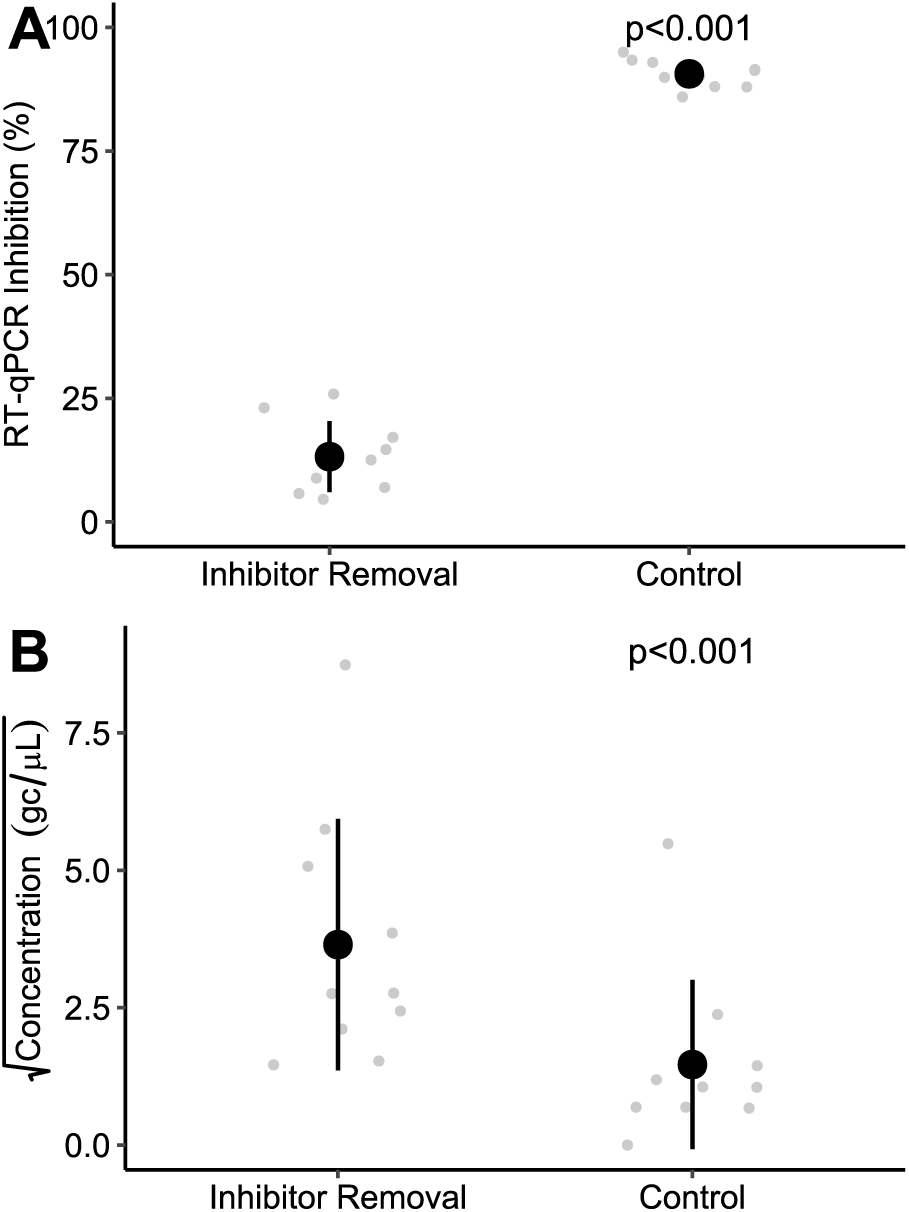
The RT-qPCR inhibition (**A**) and concentration (**B**) of norovirus GI from cDNA samples processed with or without inhibitor removal. Black markers show the average and grey show the raw data; error bars show ± 1 sd. N=10

**Figure 2.**
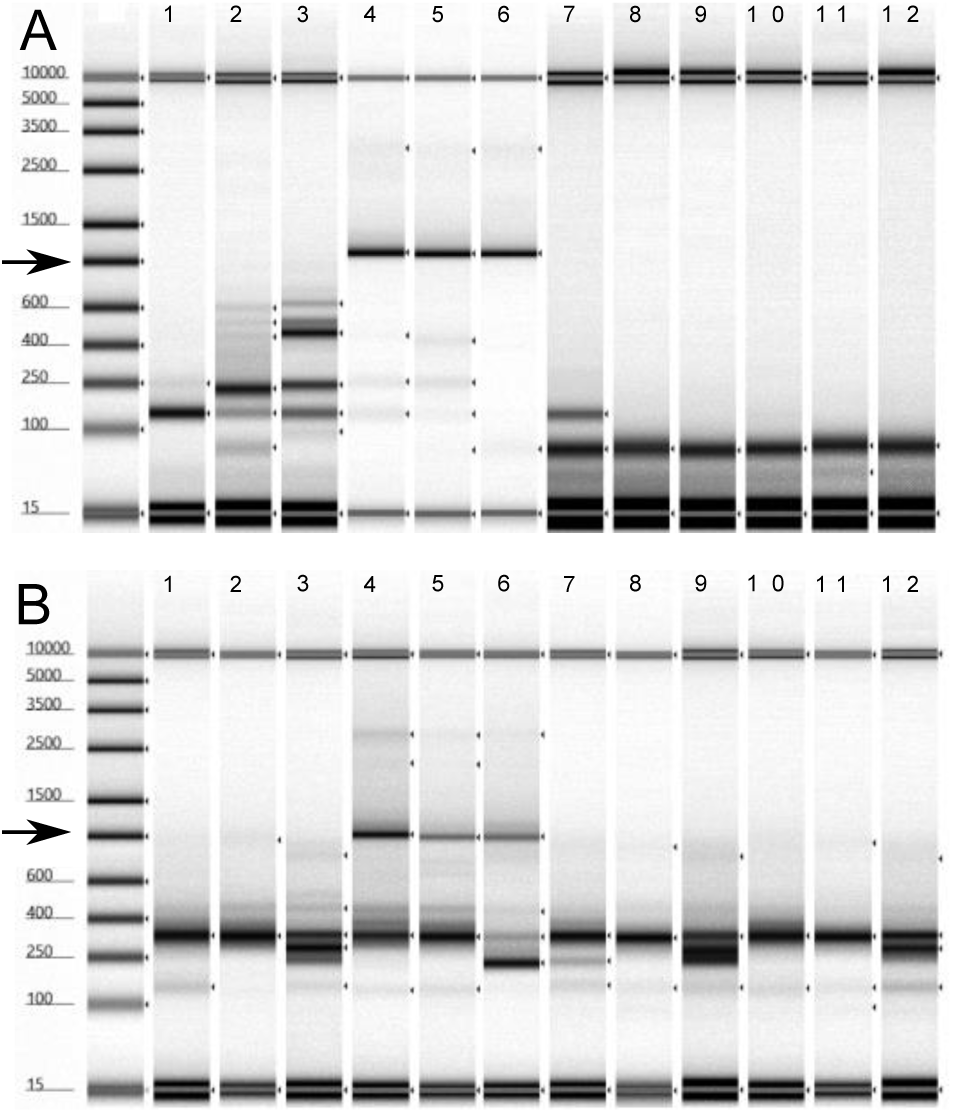
TapeStation images showing the RdRp+VP1 amplicon for GI (**A**) and GII (**B**) using two different reverse transcriptases, LunaScript® (1 to 6) and SuperScript™ (7 to 12). Samples were processed without (1 to 3 and 7 to 9) or with (4 to 6 and 10 to 12) inhibitor removal. Products are 1110 and 971 bp for GI and GII. Black arrow indicates the expected amplicon size, n= 3.

### 3.2 PCR Reagents, Cycle Optimisation and Amplicon Purification

Following first round PCR, no GI amplicons were observed indicating that the polymerases lacked the sensitivity to amplify NoV GI in a single round. For GII, however, amplicons were observed but not for all dilutions (Figure 3). Phusion™ showed signs of PCR inhibition with increased yields following dilution (Figure 3 B). Q5U® and Ultra™ II Q5® showed abundant NSA at all T_a_ (Figure 3 D and E). In general, Platinum™ and LongAmp® showed low levels of NSA, good yield and sensitivity with both showing amplification with samples diluted 10-fold at their optimal T_a_. For LongAmp™, yield reduced with increasing T_a_ but Platinum™ increased with T_a_ while SuperFi™ showed a reduction in sensitivity and NSA but an increase in yield with increasing T_a_ (Figure 3 A, C and F).

**Figure 3.**
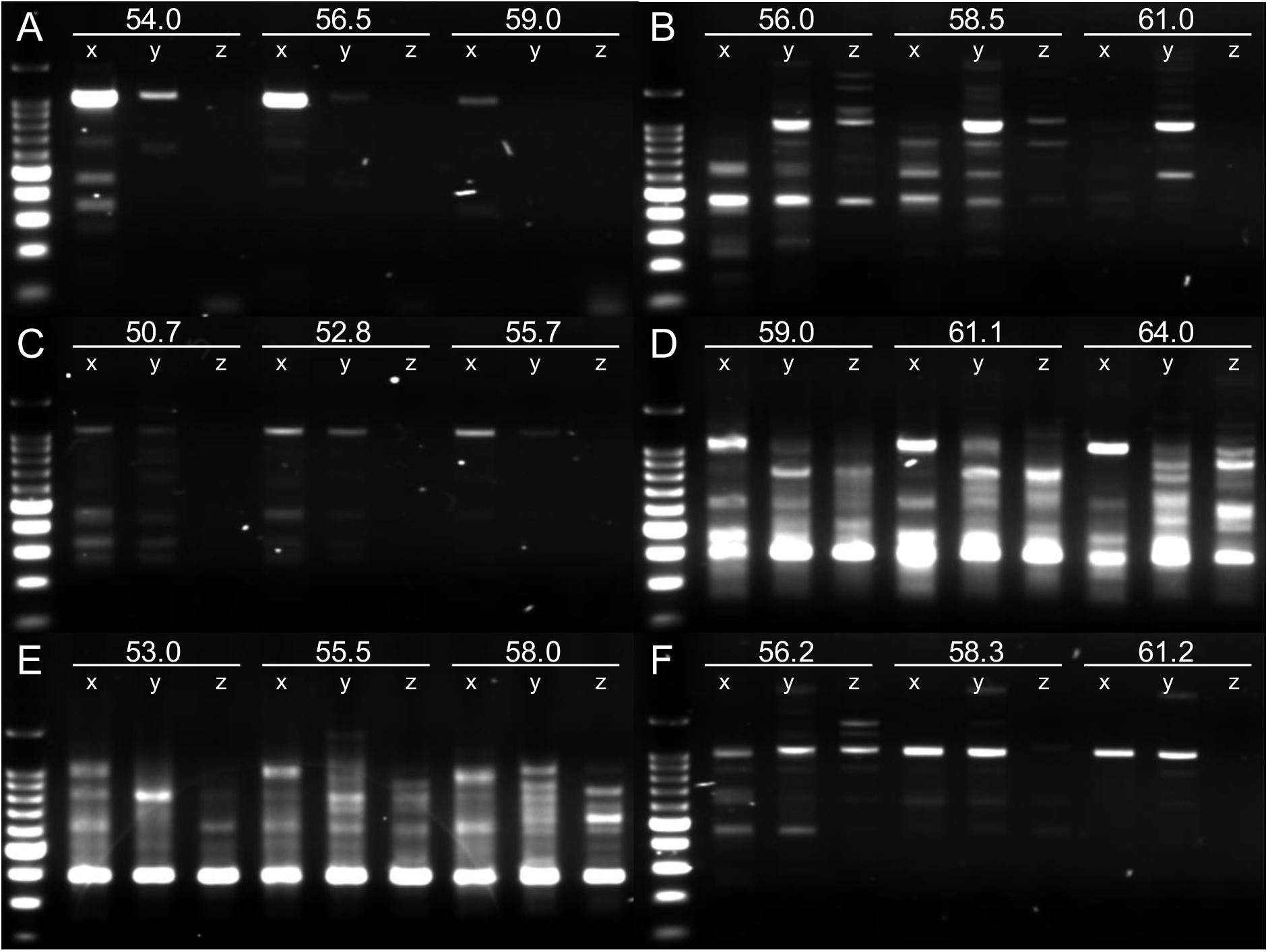
Gel electrophoresis of the first-round GII PCR products from several polymerases **A**) LongAmp®, **B**) Phusion™, **C**) Platinum™, **D**) Q5U®, **E**) Ultra™ II Q5® and **F**) SuperFi™. Numbers indicate the annealing temperature (°C) and x, y, and z are 1-, 10- and 100-fold diluted cDNA. Expected PCR products are ∼1 kb (second marker) of the 100 bp DNA ladder (Promega, USA). N=2, second sample not shown.

For the semi-nested PCR, LongAmp® and Platinum™ were pursued as SuperFi™ cannot process the inosine bases present in GISKR_m_. Platinum™ showed the best sensitivity for GI with amplification at 100-fold dilution and lower NSA compared to LongAmp® (Figure 4 A and B). Optimum T_a_ was 47.4°C and 57.2°C for the first-round and semi-nested PCR, respectively (Figure 4 A). For GII, Platinum™ showed better sensitivity compared to LongAmp® and its NSA was slightly reduced at T_a_=55.7°C for first-round and semi-nested PCR (Figure 4 C and D).

**Figure 4.**
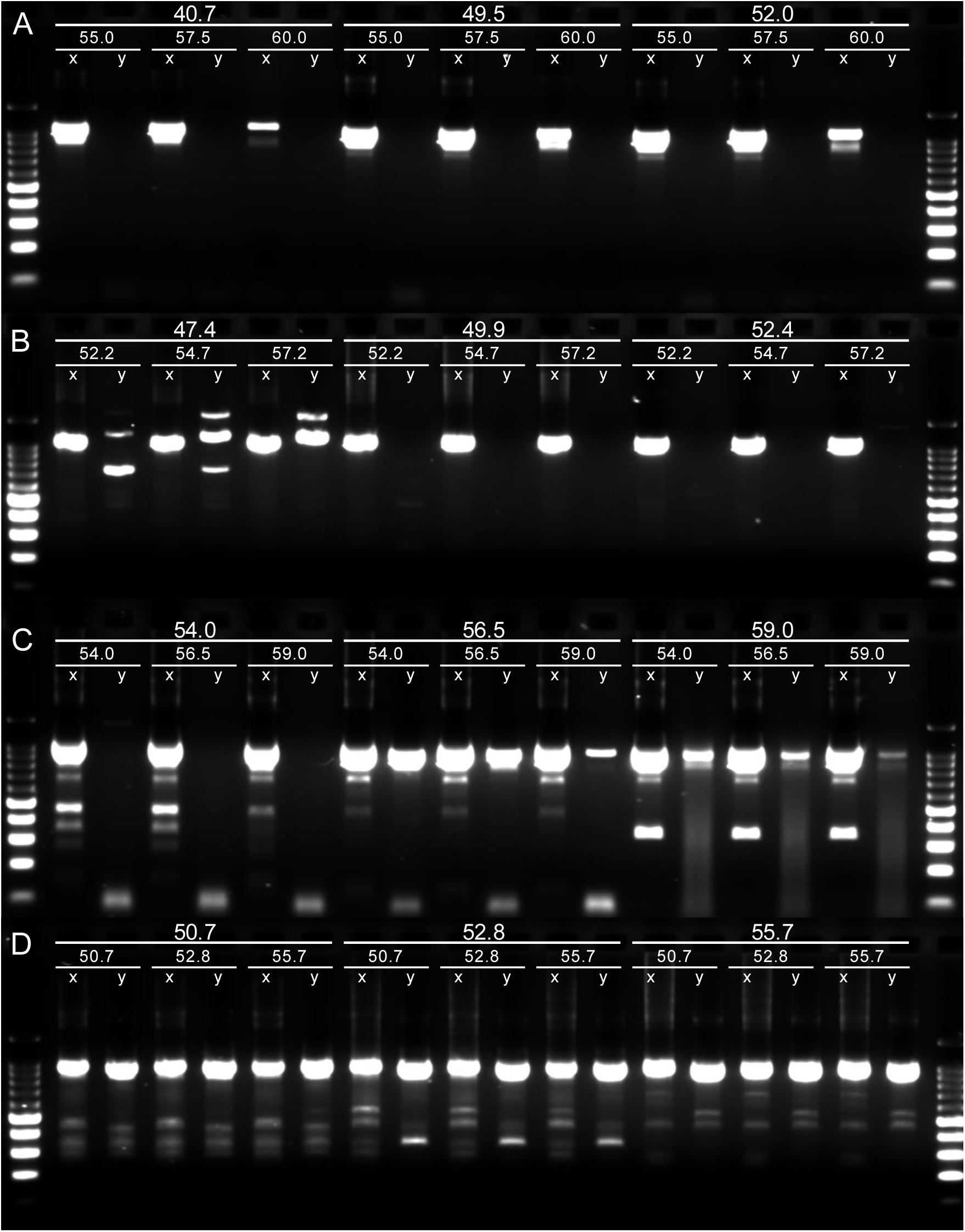
Gel electrophoresis of the semi-nested PCR for GI (**A** and **B**) and GII (**C** and **D**) using polymerases LongAmp® (**A** and **C**) and Platinum™ (**B** and **D**). Numbers indicate first-round and semi-nested annealing temperatures. Samples were run at 1 (**x**) and 100-fold (**y**) dilution. Expected PCR products are ∼1 kb (second marker) on the 100 bp DNA ladder (Promega, USA). N=2, second sample not shown.

### 3.3 Development of a Multi-Target Library

Following PCR optimisation, NSA was higher for GII than GI with the AOI% at 61.5% and 89.1% (Figure 4 B and D and Figure 5 A and B). PCR optimisation significantly increased AOI% for GI and GII from 77.6% to 89.1% and 33.4% to 61.5%(Figure 5 A and B). PCR yield significantly increased for GII (p<0.001) following optimisation from 11.84 ng/µL ± 2.71(sd) to 22.08 ng/µL ± 3.47 while GI was not impacted; 29.97 ng/µL ± 4.47 and 29.78 ng/µL ± 2.80.

**Figure 5.**
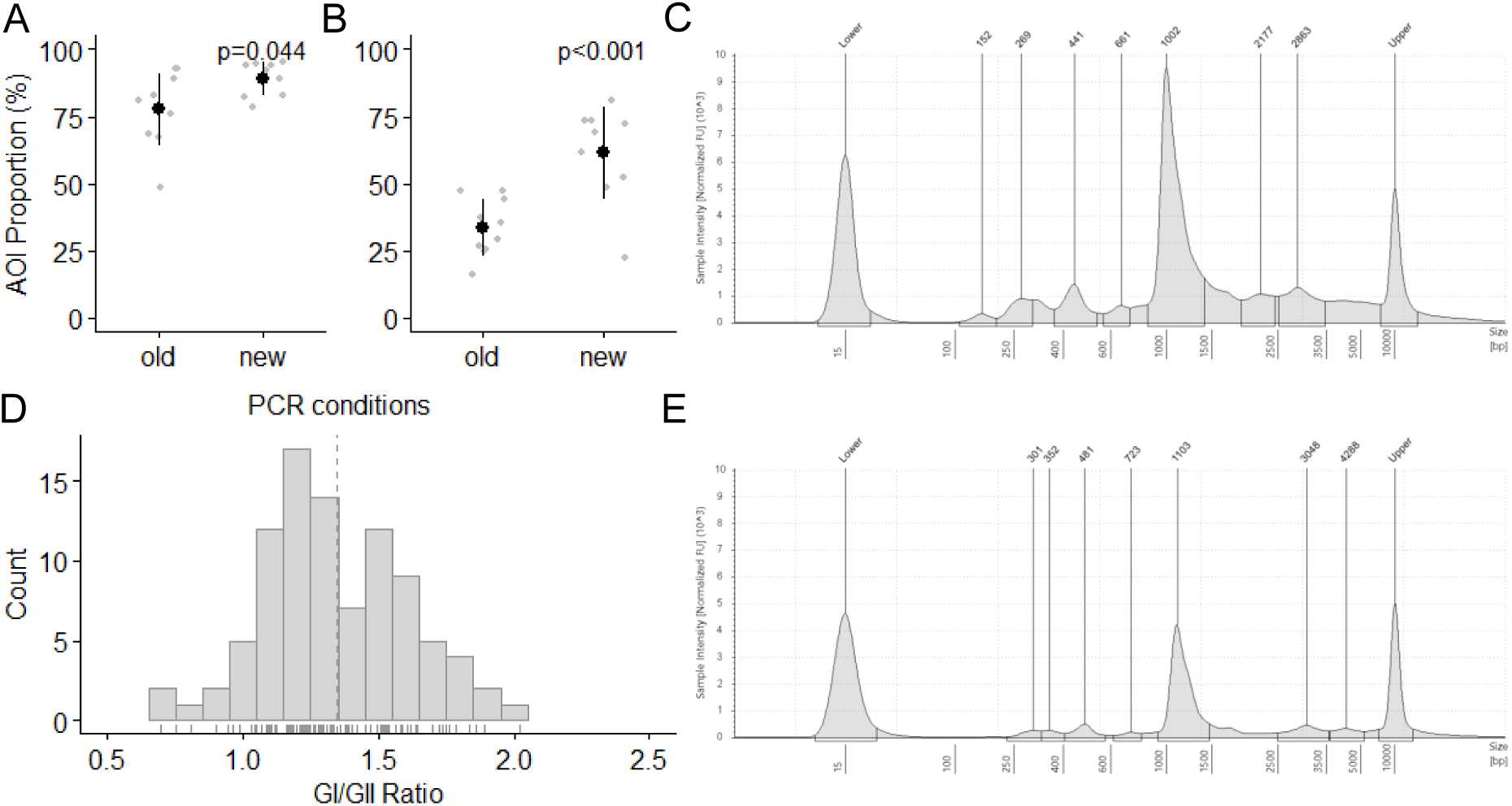
The percentage of the amplicon of interest (AOI) within the total amplicon pool for the T_a_=50°C and optimised PCR for norovirus GI (**A**) and GII (**B**). TapeStation electropherograms of norovirus GI PCR before (**C**) and after (**E**) size selection. The GI/GII AOI ratios (**D**). For **A** and **B** n = 10, the black dots show the average AOI% and grey dots show individual sample data. Error bars show ± 1 sd. For **D,** n = 130 and dashed line indicates the mean.

To reduce sequencing of NSA, Ampure XP- and Mag-Bind® TotalPure NGS-based size selection were trialled. Neither increased the AOI% (78.4 to 79.4%) but both saw reductions in amplicon concentrations (24.1 to 6.14 ng/µL); highlighted in Figure 5 C and E. A library pooling method based on the AOI% within the total amplicon pools for GI and GII was developed with a 1.34-fold mean difference between the GI and GII AOI% being identified (Figure 5 D). Adjusting the GI and GII pooling molarities reduced the bias in median read coverage from 3.3-fold to 2.6-fold.

### 3.4 Sequence Quality, Nanopore Sequence Validation and PCR Chimeras

During comparative analysis of the wastewater optimised methods and those using T_a_=50°C, whole process and PCR negative controls were free from any sequences aligning to NoV (Supplementary Table 9). Prior to reads trimming the mean and median read lengths were 1,790 and 851 bp, respectively, suggesting a positively skewed distribution with a few very long reads. Approximately 45.5% to 71.2% of reads per sample included both primers and of those 17.2% to 45.2% aligned against a consensus sequence (Supplementary Table 9). Of 94 consensus sequences, 3 couldn’t be typed due to indels in homopolymer regions.

To validate the nanopore consensus sequences, three faecal samples from assumed single-type infections were compared to sequences obtained by Sanger sequencing. All three had 100% nucleotide similarity compared to Sanger with no indels. The Sanger sequences had median and modal quality scores of Q58 (Supplementary Table 3). Alignment of nanopore reads from these samples against the consensus sequences estimated the median quality of reads in the sequencing library as being 12.3 (Supplementary Figure 1).

During initial data processing, 12 putative novel recombinant GI types were observed on 18 occasions (Supplementary Table 4). On 16 occasions, parent-types containing the genotype or p-type were both present at ≥7.4% of the total reads with the recombinants comprising ≤2.5%; indicating their potential as PCR chimeras. On two occasions the parent types (GI.1[P1] and GI.2[P2]) of the putative chimeras (GI.1[P2] and GI.2[P1]) were not detected in the sample. Some putative chimeras contained a low number of SNPs (<5) compared to their parent types.

### 3.5 The Impact of PCR Optimisation on Norovirus Diversity

Ten pooled wastewater samples were analysed to determine the impact of the PCR optimisation on norovirus taxa richness. The optimised PCR detected 13 GI types, two (GI.5[P12] and GI.7[P9]) were not detected by the T_a_=50 °C method. Following optimisation taxa observations reduced from 41 to 38 along with taxa richness from 4.1 ± 1.7 to 3.8 ± 1.4 (p=0.729) (Figure 6 i and ii). Differences in detected taxa were from types whose genotypes and p-types had been detected separately in different samples; apart from GI.P5 which was not detected by the optimised assay. For GII, the total number of types detected increased from 6 to 8 after optimisation with T_a_=50°C not detecting GII.2[P31] and GII.4[P16] (Figure 7). Total taxa count increased from 26 to 41 and taxa richness significantly increased from 2.6 ± 1.07 and 4.1 ± 0.7 (p<0.001) (Figure 7B i and ii).

**Figure 6.**
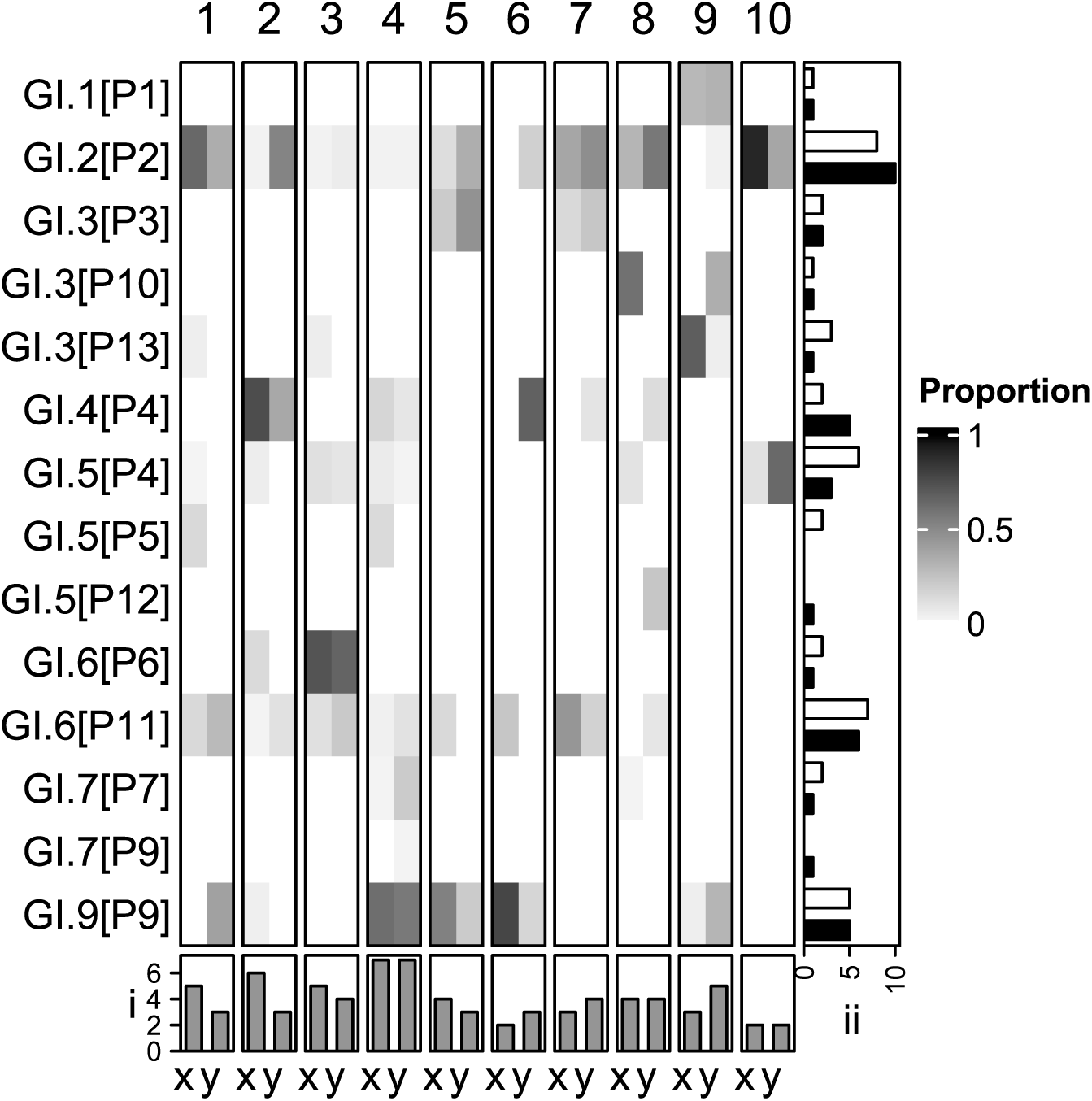
Norovirus genogroup I types identified in ten pooled wastewater samples (1 to 10) using two PCR assays (x and y). Assay x used T_a_=50°C while y is the wastewater optimised assay. The heatmap shows the proportion of reads assigned to each of the type. **i**) is taxa richness and (**ii**) is the frequency of observations for the two different methods x (white) and y (black).

**Figure 7.**
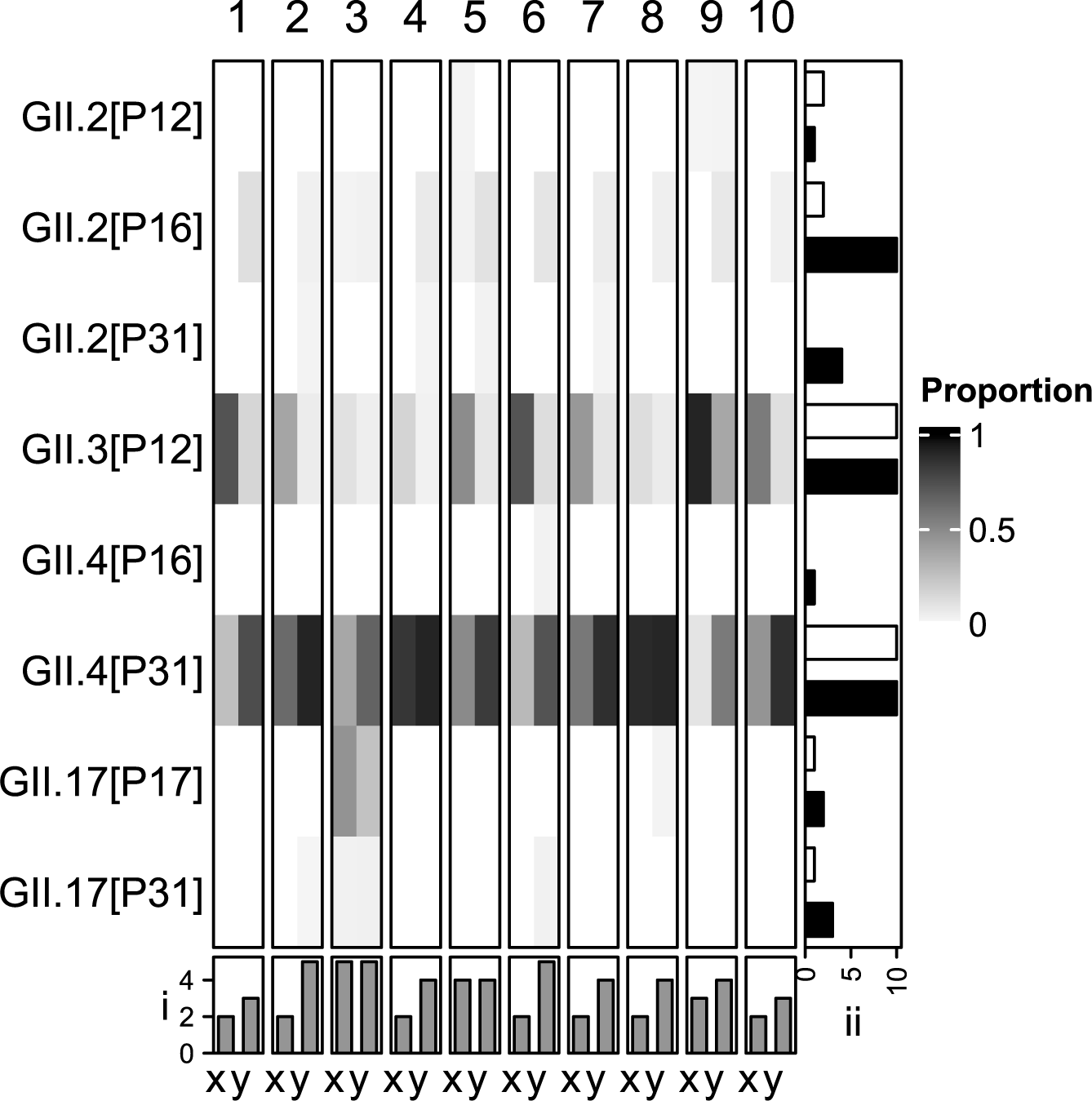
Norovirus genogroup II types identified in ten pooled wastewater samples (1 to 10) using two PCR assays (x and y). Assay x used T_a_=50°C while y is the wastewater optimised assay. The heatmap shows the proportion of reads assigned to each of the type. **i**) is taxa richness and (**ii**) is the frequency of observations for the two different methods x (white) and y (black).

### 3.6 Noroviruses in Wastewater in England

This section only uses the data from the wastewater optimised assay. Aside from GI.8, all GI genotypes and 11 out of 14 GI p-types described by the CDC were detected from the pooled samples collected from across England (Supplementary Table 5 and 6)(Centre for Disease Control, 2023b). Analysis of the genotype:p-type combinations identified up to 3 p-types per genotype and two genotypes per p-type, whilst both had a modal number of 1 pairing (Figure 6 A). Four taxa GI.2[P2], GI.4[P4], GI.6[P11] and GI.9[P9] occurred in ≥50% of the samples (100%, 50%, 60% and 50%) with GI.2[P2] and GI.9[P9] tending to be the dominant taxa when present (Figure 6).

Eight GII types, representing 4 of the 26 genotypes and 4 of the of the 37 p-types as described by the CDC were detected (Supplementary Table 7 and 8)(Centre for Disease Control, 2023b). Up to 3 p-types per genotype and 3 genotypes per p-type were observed with a mode of 2 for both. Three GII-types (GII.2[P16], GII.3[P12] and GII.4[P31]) were detected in 100% of the samples with GII.4[P31] having the highest proportion of reads in all cases (Figure 7).

## 4.0 Discussion

Two of the challenges faced during method development were the physiochemical and microbiological nature of wastewater, their impact on NoV detection and the biology of NoVs themselves. As enteric viruses, NoVs are detected frequently (82% to 100%) in wastewater making NoV negative samples difficult to obtain (Qiu *et al*., 2018; Huang *et al*., 2022). Performing method validation with spiked-matrix mock communities is, therefore, difficult to achieve. Additionally, difficulties obtaining clinical samples and culturing NoVs makes obtaining a diverse range of reference material for artificial matrix-based experimentation challenging (Cates *et al*., 2020).

Furthermore, such experiments fail to account for the potentially large matrix-effect that wastewater exhibits (Scott *et al*., 2023). This creates issues around determination of PCR chimeras, quantitative validation and calculating sequencing error rates.

### 4.1 Wastewater and RT-PCR Inhibition

Wastewater’s physicochemical properties are poorly characterised and are likely influenced by many geospatial and environmental factors. RT and PCR inhibitors likely present include polysaccharides, bile salts, lipid, urate, fulvic and humic acids, metal ions, algae and polyphenols (Schrader *et al*., 2012; Sims and Kasprzyk-Hordern, 2020). Inter-sample RT-qPCR inhibition levels have been shown to be highly variable (0% to 98%) for SARS-CoV-2 (Scott *et al*., 2023). Inhibition is, therefore, likely to influence the data quality of PCR-dependent WBE methods.

Here, we confirm the suitability of Child *et al*., (2023)’s inhibitor removal method for detecting NoVs in wastewater, reducing inhibition (85%) and increasing NoV quantification (4.8-fold) as measured by RT-qPCR (Figure 1 A and B). These data should be used with caution as inhibitor susceptibility may change between different enzymes and amplicons; quantitative performance gains for this metabarcoding approach cannot be inferred (Huggett *et al*., 2008; Kermekchiev *et al*., 2009). Following inhibitor removal, LunaScript™ showed 163- to 1731-fold increases in sequencing depth highlighting the large impact inhibitors can have on amplicon sequencing and the importance of matrix-specific assay optimisation.

### 4.2 Wastewater and PCR Optimisation

The importance of matrix-specific assay optimisation is emphasised by wastewater’s likely diverse nucleic acid content with influence from humans, their food, microbial ecology of the gastrointestinal tract other natural and industrial sources of wastewater. This makes designing nucleic acid-based assays challenging due to an increased likelihood of NSA. Furthermore, nucleic acid degeneracy within primers is required to capture the full genomic diversity of NoV; increasing the likelihood of NSA (De Graaf et al., 2016; Ford-Siltz et al., 2021).

Several polymerases were assessed for sensitivity, NSA and yield. Platinum™ showed the best performance overall for both GI and GII PCR assays (Figure 3 & Figure 4). It is likely that Platinum™ shows increased enzymatic activity at higher temperatures allowing good performance at higher T_a_ thereby reducing NSA and increasing yields. Optimisation of the Platinum™ PCR conditions significantly increased taxa richness for GII (57.7%) while a non-significant reduction was seen for GI. Assay optimisation increased the frequency of observation for some taxa; for example GII.2[P16] was detected in 20% of the samples using the T_a_=50°C method but in 100% with the optimised protocol. Others (GII.2[P31] & GII.4[P16]) weren’t detected using the T_a_=50°C method (Figure 6 B). This is likely due increased T_a_ reducing primer-template interactions when mismatches are present allowing the preferential amplification of NoV. Failure to properly optimise methods implemented in WBE is, therefore, likely to underestimate the diversity of the organisms under investigation.

### 4.3 Sequencing and Bioinformatics

GI and GII were sequenced under a single barcode per sample to maximise throughput. PCRs were optimised to reduce NSA but the residual NSA led to read-depth disparity for GI and GII. Size selection to remove NSA was unsuccessful, perhaps because products of the NSA were often close in size to the AOI. A weighting was therefore applied to the loading of the GI and GII amplicons into library prep to account for the differences in depth.

Given the high estimated read error rate (median ≈Q13) a consensus-based approach was used and analysis was focused mainly at the type-level. Consensus sequences were clustered at 95% prior to estimating abundance of the different sequences within the dataset. This threshold allowed for differentiation of types while accounting for the significant amount of noise in the data. GII sub-types, however, cannot be determined as heterogeneity thresholds are 98% (Tatusov *et al*., 2021). Consensus sequences used to identify novel recombinants and putative PCR chimers were supported by high-quality alignments of ≥900 bp in length, included ≥90% of a read and had normalised read scores of ≥92%.

A large amount of sequencing data were filtered prior to alignment, which appeared to be due to NSA and the high error rate inherent in nanopore sequencing. The latter should improve with new developments in nanopore sequencing chemistry, basecalling and flowcells. Despite most of the analysis focussing on types, rather than subtypes, it was shown when analysing NoV faecal samples from three presumed single type infections there was 100% nucleotide similarity with Sanger sequencing; with the latter having a median basecalling score of Q58 (Supplementary Table 3). This indicates that this method is adequate to discriminate between subtypes if the consensus sequence clustering is performed at 98% rather than the 95% implemented here.

### 4.4 PCR Chimeras

Twelve putative novel GI *RdRp+VP1* recombinants were identified but removed as putative PCR chimeras. For most cases, putative chimeras were detected alongside parent types at higher abundance. On two occasions, only a single parent was identified in the same sample, increasing the likelihood that identification of a novel recombinant was genuine. Chimera formation without the detection of the parent, however, has previously been observed by Ollivier *et al*., (2022).

Following artificial bioaccumulation of oysters with GII.4[P16], GII.2[P16], GII.4[P31] and GII.17[P17] the detection of two chimeric types GII.17[P16] and GII.17[P13] occurred without detection of the parent type GII.17[P17]. This method used a shorter amplicon (426 bp) targeting the ORF-1 and -2 junction using Illumina MiSeq. Failure to detect both parent types may indicate failure of the forward primers to amplify GII.P17. In this study, however, the two parent types (GI.1[P1] and GI.2[P2]) of the putative chimeras (GI.1[P2] and GI.2[P1]) have been successfully detected in other samples (Figure 6). This gives further indication that the novel recombinants may exist in nature.

Both putative chimeras, however, formed a low proportion (0.4%) of the total reads (Supplementary Table 4). Due to wastewaters high probability of containing degraded nucleic acids, further investigation is required to determine whether these novel recombinants are genuine. Degradation of nucleic acids may prevent the detection of a parent type but still allow it to form PCR chimeras at low concentrations. This increases the likelihood of chimera formation from that of prematurely terminated strand elongation alone (Meyerhans et al., 1990). These putative chimeras were removed from the data due to the lack of additional evidence that they are genuine. Some putative chimeras contained up to 5 SNPs difference from the parent types. Although SNP presence usually provides evidence against PCR chimera formation, these SNPs may have been introduced through the clustering of consensus sequences at 95% and as such the sequences were also removed.

Differentiating between true recombinants and PCR chimeras *in-silico* is challenging. Reference-based chimera filtering methods will fail to detect novel recombinants or remove chimeras formed from known types. De-novo approaches rely on read depth to determine parent and chimeric sequences and may potentially including chimeric sequences with similar read depths or skews in read depth caused by PCR bias. This is highlighted by the USEARCH de-novo approach missing several potential chimeras which were removed following manual screening. Additional investigation into the natural NoV recombination breakpoints did not assist in identification of chimeras due to its variability within the terminal and proximal ends of *RdRp* and *VP1* (Bull *et al*., 2005; Fu *et al*., 2019).

PCR chimera formation, however, is not an issue isolated to long amplicon sequencing methods. Putative PCR chimeras were identified by Mabasa *et al*., (2022) who detected with 51 putative novel recombinants using a shorter amplicon (≈575 bp). Although shorter amplicons were used, PCR chimera formation cannot be discounted as their formation will be influenced by polymerase choice, PCR conditions and sample nucleic acid diversity (Nagai *et al*., 2022). Future research should focus on methods for confirming novel recombinants such as shorter, specific RT-PCR companion assays with low-frequency chimera production. Due to the difficulties discriminating PCR chimeras from genuine recombinants and in the absence of readily available reference material for mock community analysis to set PCR chimera filtering thresholds, it may be appropriate for metagenomic approaches to be used for the identification of novel recombinants.

### 4.5 Noroviruses in Wastewater in England

The GI.2 genotype (GI.2[P2]) was detected in 100% of samples in this study, this is in contrast to the Office for National Statistics’ (ONS) reported clinical data for the same period which stated GI.3 and GI.6 as the most frequently detected genotypes (3%) across the year; no p-type data were available (Office for National Statistics, 2023). This discrepancy, however, may be explained by the absence of GI.2 in clinical cases due to its predominantly asymptomatic infections. GI.6 (GI.6[P11] and GI.6[P6]) were found in 60% of the samples while GI.3 types (GI.3[P3], GI.3[P10] and GI.3[P13]) present in 30% supporting their reporting in the clinical data (Figure 6). Our methods, however, also detected GI.9, GI.4 and GI.5 in over 40% of the samples indicating that clinical data may underestimate NoV diversity within the population.

Greater congruence with the ONS data was seen for GII with three GII-types (GII.2[P16], GII.3[P12] and GII.4[P31]) detected in 100% of the samples and GII.4[P31] having the highest proportion of reads in all cases. The ONS reported 48% of cases were GII.4, 13% were GII.3 and 10% were GII.2 (Office for National Statistics, 2023)(Figure 7). Without further validation, however, using these data in a quantitative fashion should be done so with caution.

Comparing these two data sets is problematic as the ONS data are a summary of clinical cases in England across the year whereas the data from the present study are a snapshot of the norovirus diversity across England over several days. Differences between the two data sets, especially for GI, may also be due to the low case numbers reported (45) compared to GII (514) (Office for National Statistics, 2023). Furthermore, the samples used in this study were pooled, and any highly localised outbreaks could be out of detectable range. Time-course studies with greater geospatial separation need to be conducted to assess the link between WBE and clinical data.

## 5.0 Conclusions

We have developed a long amplicon sequencing method using nanopore sequencing that allows the comprehensive (genotyping and p-typing) and simultaneous typing of NoV GI and GII in wastewater. RT, DNA polymerase, PCR conditions and library-pooling were all optimised along with the development of a consensus-based bioinformatics pipeline. We have shown that failure to properly optimise assays for implementation in WBE is likely to lead to underestimation of taxa-richness through loss of data to NSA. Furthermore, the appropriate bioinformatic interventions are required to prevent the reporting of data from PCR chimeras.

Partial qualitative validation has been performed with 88.9% and 78.6% of GI genotypes and p-types and 15.4% and 10.8% of the GII genotypes and p-types detected. Further validation is recommended before adoption at scale. Initial analysis indicated that our data matched the GII clinical observations reported by the ONS although comparisons were difficult due to the nature of the data and lack of p-type reported in the clinical data. Future studies should focus on building a collection of NoV reference material and NoV negative wastewater. This would allow for quantitative and qualitative assay validation and optimisation of PCR methods to minimise PCR chimeras. The development of methods for confirming novel NoV recombinants should also be prioritised.

Deploying such a technique into a WBE scheme paired with additional studies into NoV shedding rates and virulence would allow better understanding of NoV diversity and prevalence within the population, outbreak tracking and insights into genetic diversity and the importance of recombination in NoV evolution. This increased understanding of the biology and epidemiology of noroviruses could lead to improvements in disease prevention and management and predictions of economic burden.

## Supporting information

Supplementary Protocol

## 6.0 Acknowledgements

First, we’d also like to acknowledge all the staff at Cefas involved in the co-ordination and management of PATH-SAFE. Second, we’d like to recognise the extraordinary effort from the members of the Environmental Monitoring for Health Protection programme for implementing and undertaking SARS-CoV-2 WBE across England from which the samples used in this study were taken. Finally, special thanks are directed to the teams involved in wastewater sample collection and all the staff at the Environment Agency’s National Laboratory Service who were involved in daily sample processing as this study couldn’t have been performed without them.

## Author Contributions

Conceptualisation: Batista, F., Lowther, J. & Scott, G.

Methodology: Batista, F., Scott, G., Ryder, D. & Treagus, S.

Investigation: Scott, G., Ryder, D. Buckley, M., Hill, R., Batista, F., Stapleton, T. & Treagus, S.

Formal Analysis: Scott, G. & Ryder, D.

Visualisation: Scott, G. & Ryder, D.

Writing – original draft: Scott, G.

Writing – review and editing: Scott, G., Batista, F, Ryder, D. Walker, D. I. & Lowther, J.

Supervision: Batista, F.

## Funding

Funded by His Majesty’s Treasury Shared Outcome Fund PATH-SAFE programme.

## Conflicts of Interest

There are no conflicts of interest to declare.

## Data Accessibility

Raw sequencing data comparing the wastewater optimised and unoptimised methods has been deposited to the National Center for Biotechnology Information databases under BioProject PRJNA1087315. Raw nanopore data and sanger sequences from the norovirus positive human faecal material is available at the BioProject and GenBank under the accessions PP542636, PP542646 and PP542648. The methods developed in this study are available as a supplementary protocol or on protocols.io dx.doi.org/10.17504/protocols.io.8epv5xpmjg1b/v1.

## Notes

https://www.ncbi.nlm.nih.gov/bioproject/PRJNA1087315

### Competing Interest Statement

The authors have declared no competing interest.

