## Supplementary Materials for "Long Amplicon Nanopore Sequencing for Dual-Typing *RdRp* and *VP1* Genes of Norovirus Genogroups I and II in Wastewater"

### 1.0 Supplementary Figures

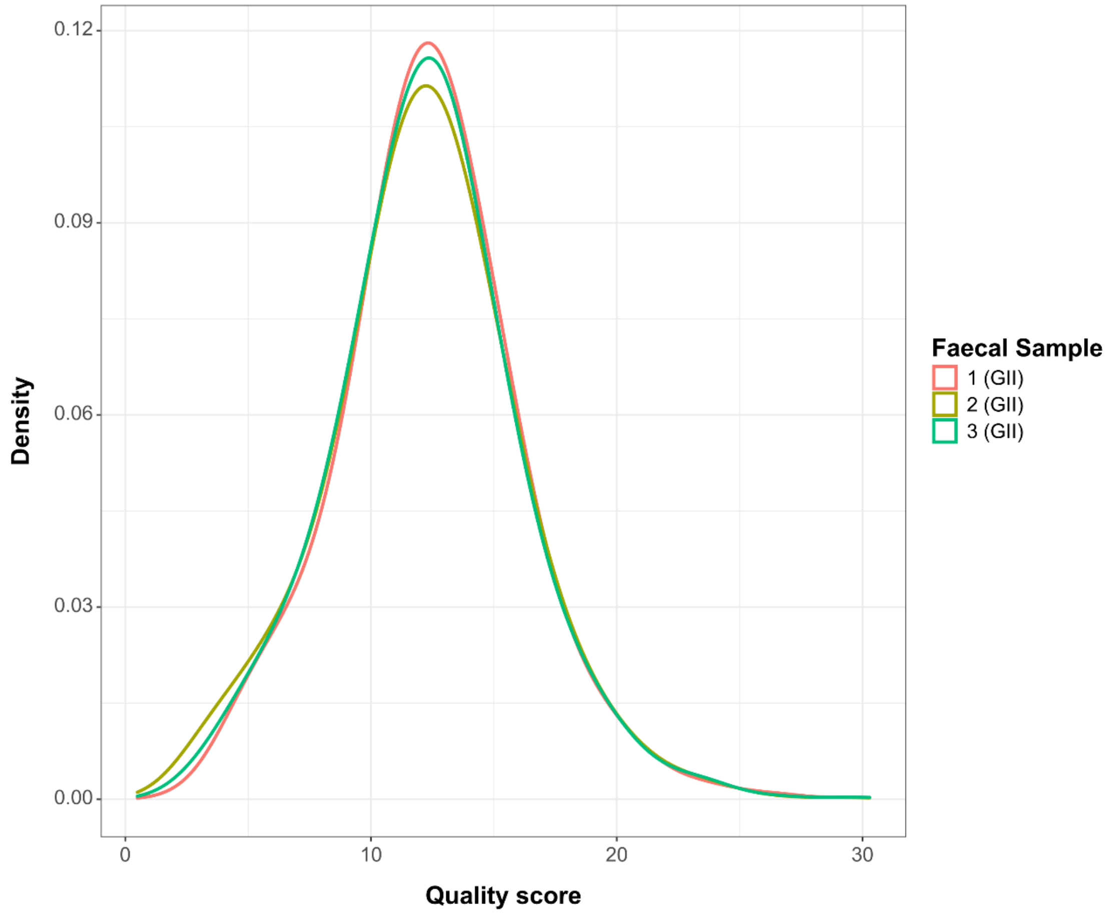

*Supplementary Figure 1: The estimated quality of reads in three faecal samples assumed be associated with single-type infections.*

### 2.0 Supplementary Equations

$$P = 100\frac{m_a}{\sum_{i=1}^{n} m}$$

*Supplementary Equation 1. Calculating the percentage the amplicon of interest (P) within the total amplicon pool where $m_a$ is the molarity of the amplicon of interest and m is the molarity of the different PCR products within a given sample.*

$$R = \frac{P_{GI}}{P_{GII}}$$

*Supplementary Equation 2. The ratio of the percentage of the amplicon of interest (R) between the norovirus genogroups I and II PCRs. P is as calculated in Equation 1 for genogroups I and II.*

$$GI\ moles = \frac{200}{R+1}$$

*Supplementary Equation 3. The moles of GI PCR product in fmol (GI moles) to be input into end-prep where R is the ratio of GI and GII amplicons as calculated in Equation 2.*

$$GII\ moles = R\frac{200}{R+1}$$

*Supplementary Equation 4. The moles of GII PCR product in fmol (GII moles) to be input into end-prep where R is the ratio of GI and GII amplicons as calculated in Equation 2.*

$$p = \frac{L_m - L_o}{L_o}$$

$$Q = -10\, log_{10}\, p$$

*Supplementary Equation 5. Estimating the probability that a base call is incorrect (p) and the overall quality of reads (Q) using the number of matching bases ($L_m$) and the overall length of the alignment ($L_o$) reported by minimap2.*

### 3.0 Supplementary Tables

*Supplementary Table 1. The pooled samples used in this study, their associated regions and the number of nucleic acid extracts in each composite.*

| Sample | Regions | No. of samples |
|---|---|---|
| 1 | East and West Midlands | 34 |
| 2 | East of England | 29 |
| 3 | North West | 28 |
| 4 | South East | 14 |
| 5 | South West | 14 |
| 6 | Yorkshire | 24 |
| 7 | South East | 14 |
| 8 | South East | 21 |
| 9 | South West | 14 |
| 10 | South West | 17 |

*Supplementary Table 2. Bioinformatic procedures, tools and commands.*

| Procedure | Tool | Parameters | Reference |
|---|---|---|---|
| Basecalling and de-multiplexing | Guppy 6.3.9 | Flow cell type 'FLO-MIN114' Kit type 'SQK-NBD114-96' | Oxford Nanopore Technologies, 2024 |
| Read splitting | duplex_tools 0.2.14 | --allow_multiple_splits Native | Oxford Nanopore Technologies, 2022 |
| Read trimming | cutadapt 3.4 | --action=trim -n 1 --discard-untrimmed -e 0.30 -O 12 –revcomp -g file:primer_sequences.fasta | Martin, 2011 |
| Size filtering | SeqTK 1.3 | seq -L 800 | Li, 2018 |
| Random sampling | SeqTK 1.3 | sample 90000 | Li, 2018 |
| Identifying read overlaps | Minimap 2.24 | -k19 -Xw19 -e0 -m100 -r100 -I 30M --cap-kalloc=8000m --cap-sw-mem=100m | Heng Li, 2018 |
| Identification of chimeras and sequences with poor support | yacrd 1.0.0 | -c 10 -n 0.2 | Marijon, Chikhi and Varré, 2020 |
| Read clustering and consensus sequence generation | NGSpeciesID 0.1.3 | --q 15 --ont --consensus --max_seqs_for_consensus 10000 --rc_identity_threshold 0.9 --aligned_threshold 0.8 --mapped_threshold 0.8 --m 1100 --s 200 | Sahlin, Lim and Prost, 2021 |
| Indexing of consensus sequences | kma 1.4.9 | -NI | Clausen, Aarestrup and Lund, 2018 |
| Alignment, variant calling, and masking of poorly supported regions | kma 1.4.9 | -vcf 1 -sam -ConClave 2 -bcNano -bc 0.7 -bcd 100 -t 20 -md 100 -1t1 -ont -ml 900 -xl 1100 -ef -mrs 0.92 -mrc 0.90 | Clausen, Aarestrup and Lund, 2018 |
| Rename consensus sequences | SeqKit 2.3.0 | rename sequences.fasta OTU_ | Shen *et al.*, 2016 |
| Indexing of consensus sequences | Samtools 1.13 | -faidx | Danecek *et al.*, 2021 |
| Tabulating regions which were identified and masked by kma | SeqKit 2.3.0 | locate --bed -P -r -p '^[agct]+' -p '[agct]+$' | Shen *et al.*, 2016 |
| Sorting and complementing coordinates, and using the result to trim seqs | Bedtools 2.30.0 | The sort, complement and getfasta commands were applied using output from the SeqKit and Samtools command. | Quinlan and Hall, 2010 |
| Clustering of consensus sequences | CD-HIT 4.8.1 | cd-hit-est -G 0 -c 0.95 -n 10 -d 0 -M 100000 -T 12 -g 1 -aL 0.9 | Fu *et al.*, 2012 |
| Removal of PCR chimeras | USEARCH 11 | -uchime3_denovo -chimeras | Edgar, 2016 |

*Supplementary Table 3.* Nanopore sequence validation against Sanger sequencing.

| Faecal Sample | Genogroup | Genotype and Subtype | Nanopore length (bp) | Sanger length (bp) | Q30 Sanger length (bp) | Indels | Nucleotide similarity (%) |
|---|---|---|---|---|---|---|---|
| 1 | GII | GII.4 New Orleans [P4] New Orleans | 1008 | 945 | 862 | 0 | 100 |
| 2 | GII | GII.4 Sydney [P31] | 977 | 926 | 875 | 0 | 100 |
| 3 | GII | GII.4 Sydney [P31] | 926 | 951 | 884 | 0 | 100 |

*Supplementary Table 4. Novel norovirus GI types detected which may be PCR chimeras.*

| Type | Sample | Read proportion (%) | Genotype parent read proportion (%) | P-Type parent read proportion (%) |
|---|---|---|---|---|
| GI.1[P2] | 5 | 0.4 | 0 | 33 |
| GI.2[P11] | 2 | 0.8 | 52 | 27 |
| GI.2[P11] | 7 | 2.5 | 47 | 17 |
| GI.2[P12] | 8 | 1.1 | 57 | 23 |
| GI.2[P1] | 5 | 0.4 | 33 | 0 |
| GI.2[P3] | 7 | 0.4 | 47 | 22 |
| GI.2[P4] | 2 | 1.2 | 52 | 36 |
| GI.2[P4] | 6 | 0.7 | 18 | 67 |
| GI.2[P4] | 7 | 0.4 | 47 | 8 |
| GI.4[P11] | 4 | 0.8 | 7 | 9 |
| GI.4[P2] | 2 | 0.8 | 36 | 52 |
| GI.6[P2] | 7 | 1.0 | 17 | 47 |
| GI.6[P3] | 7 | 0.6 | 17 | 22 |
| GI.6[P4] | 3 | 0.9 | 21 | 8 |
| GI.6[P4] | 4 | 0.6 | 9 | 7 |
| GI.6[P4] | 7 | 0.4 | 17 | 8 |
| GI.9[P3] | 5 | 0.6 | 20 | 46 |
| GI.9[P7] | 4 | 1.0 | 57 | 19 |

*Supplementary Table 5. Norovirus GI genotypes detected.*

| Genotype | Detected |
|---|---|
| GI.1 | Yes |
| GI.2 | Yes |
| GI.3 | Yes |
| GI.4 | Yes |
| GI.5 | Yes |
| GI.6 | Yes |
| GI.7 | Yes |
| GI.8 | No |
| GI.9 | Yes |

*Supplementary Table 6. Norovirus GI polymerase types detected.*

| Polymerase type | Detected |
|---|---|
| GI.P1 | Yes |
| GI.P2 | Yes |
| GI.P3 | Yes |
| GI.P4 | Yes |
| GI.P5 | No |
| GI.P6 | Yes |
| GI.P7 | Yes |
| GI.P8 | No |
| GI.P9 | Yes |
| GI.P10 | Yes |
| GI.P11 | Yes |
| GI.P12 | Yes |
| GI.P13 | Yes |
| GI.P14 | No |

*Supplementary Table 7. Norovirus GII genotypes detected.*

| Genotype | Detected |
|---|---|
| GII.1 | No |
| GII.2 | Yes |
| GII.3 | Yes |
| GII.4 | Yes |
| GII.5 | No |
| GII.6 | No |
| GII.7 | No |
| GII.8 | No |
| GII.9 | No |
| GII.10 | No |
| GII.11 | No |
| GII.12 | No |
| GII.13 | No |
| GII.14 | No |
| GII.16 | No |
| GII.17 | Yes |
| GII.18 | No |
| GII.19 | No |
| GII.20 | No |
| GII.21 | No |
| GII.22 | No |
| GII.23 | No |
| GII.24 | No |
| GII.25 | No |
| GII.26 | No |
| GII.27 | No |

*Supplementary Table 8. Norovirus GII polymerase types detected.*

| Polymerase type | Detected |
|---|---|
| GII.P1 | No |
| GII.P2 | No |
| GII.P3 | No |
| GII.P4 | No |
| GII.P5 | No |
| GII.P6 | No |
| GII.P7 | No |
| GII.P8 | No |
| GII.P11 | No |
| GII.P12 | Yes |
| GII.P13 | No |
| GII.P15 | No |
| GII.P16 | Yes |
| GII.P17 | Yes |
| GII.P18 | No |
| GII.P20 | No |
| GII.P21 | No |
| GII.P22 | No |
| GII.P23 | No |
| GII.P24 | No |
| GII.P25 | No |
| GII.P26 | No |
| GII.P27 | No |
| GII.P28 | No |
| GII.P29 | No |
| GII.P30 | No |
| GII.P31 | Yes |
| GII.P32 | No |
| GII.P33 | No |
| GII.P34 | No |
| GII.P35 | No |
| GII.P36 | No |
| GII.P37 | No |
| GII.P38 | No |
| GII.P39 | No |
| GII.P40 | No |
| GII.P41 | No |

Supplementary Table 9. Trimming statistics from the nanopore sequencing run

| Sample | PCR | Prior to trimming | After trimming | Aligned |
|---|---|---|---|---|
| 1 | Optimised | 189871 | 115837 | 25478 |
| 1 | Ta=50 | 343517 | 178910 | 35109 |
| 2 | Optimised | 296867 | 186637 | 66655 |
| 2 | Ta=50 | 269260 | 151823 | 53328 |
| 3 | Optimised | 101838 | 72567 | 28005 |
| 3 | Ta=50 | 228617 | 138937 | 39936 |
| 4 | Optimised | 211179 | 141071 | 46934 |
| 4 | Ta=50 | 244136 | 137252 | 41135 |
| 5 | Optimised | 176652 | 120018 | 50043 |
| 5 | Ta=50 | 202231 | 114026 | 34378 |
| 6 | Optimised | 127933 | 77030 | 33453 |
| 6 | Ta=50 | 160346 | 76639 | 29239 |
| 7 | Optimised | 239487 | 153016 | 53502 |
| 7 | Ta=50 | 215035 | 97940 | 16830 |
| 8 | Optimised | 127411 | 82976 | 27813 |
| 8 | Ta=50 | 218580 | 127103 | 42306 |
| 9 | Optimised | 118572 | 77795 | 35158 |
| 9 | Ta=50 | 314565 | 181594 | 45298 |
| 10 | Optimised | 80460 | 50634 | 22486 |
| 10 | Ta=50 | 193210 | 99241 | 36497 |
| Replicate 1 process control | Optimised | 1463 | 484 | 0 |
| Replicate 1 process control | Ta=50 | 1150 | 460 | 0 |
| Replicate 2 process control | Optimised | 1391 | 422 | 0 |
| Replicate 2 process control | Ta=50 | 901 | 393 | 0 |
| Replicate 3 process control | Optimised | 837 | 325 | 0 |
| Replicate 3 process control | Ta=50 | 1558 | 457 | 0 |
| GI PCR 1 control | Optimised | 936 | 313 | 0 |
| GI PCR 1 control | Ta=50 | 1042 | 411 | 0 |
| GI PCR 2 control | Optimised | 1456 | 482 | 0 |
| GI PCR 2 control | Ta=50 | 1145 | 434 | 0 |
| GII PCR 1 control | Optimised | 742 | 260 | 0 |
| GII PCR 1 control | Ta=50 | 1403 | 545 | 0 |
| GII PCR 2 control | Optimised | 878 | 318 | 0 |
| GII PCR 2 control | Ta=50 | 1321 | 523 | 0 |
