## Supplementary Protocol for "Long Amplicon Nanopore Sequencing for Dual-Typing *RdRp* and *VP1* Genes of Norovirus Genogroups I and II in Wastewater"

Available at protocols.io: dx.doi.org/10.17504/protocols.io.8epv5xpmjg1b/v1

### 1.0 Introduction

This protocol outlines the procedures for dual-typing (genotyping and polymerase typing) of norovirus genogroups I and II (GI and GII) from wastewater. It assumes that wastewater sample collection, viral concentration and nucleic acid extraction has already been performed. The wastewater samples used during method development were processed using an ammonium sulphate precipitation (150 mL sample) followed by nucleic acid extraction using Kingfisher Flex™ (Thermo Scientific™, UK) and NucliSENS® reagents (BioMérieux, France).

Adapting this technique for use with differing viral concentration methods and nucleic acid extraction techniques should still be suitable. The analysis of different sample matrices will also likely be possible following sample processing (up to and including nucleic acid extraction) with methods appropriate for that matrix.

This protocol starts with reverse transcription followed by semi-nested PCR amplifying the *RdRp+VP1* region of norovirus GI and GII independently. GI and GII amplicons are combined into a single library for simultaneous sequencing of the ≈1000 bp PCR products using either the Oxford Nanopore Technologies MinION or GridION. Consensus sequences are generated using NGSpeciedID then grouped at 95%. Consensus sequences are generated using NGSpeciedID, grouped at 95% then dual-typed giving both genotype and polymerase type (e.g.GI.2[P2]). Reads are aligned to consensus sequences and PCR chimeras are filtered using reference, de novo and manual approaches.

### 2.0 Materials

*Table 1. The materials required to perform the optimised protocol for dual-type sequencing of norovirus genogroups I and II from wastewater*

| Materials | Manufacturer | Product code |
|---|---|---|
| Mag-Bind® TotalPure NGS | OmegaBIO-TEK | M1378-00 |
| Ethanol, Molecular Biology Grade | Sigma-Aldrich | 1085430250 |
| Water, Molecular Biology Grade | Sigma-Aldrich | W4502-1L |
| LunaScript® RT SuperMix Kit | New England Biolabs | E3010 |
| TE buffer, Molecular Biology Grade | Sigma-Aldrich | 574793 |
| Platinum™ Taq Polymerase | Invitrogen | 10966018 |
| dNTP Deoxynucleotide (dNTP) Solution Mix | New England Biolabs | N0447S |
| D5000 Reagents, TapeStation | Agilent | 5067-5589 |
| D5000 ScreenTapes, Tapestation | Agilent | 5067-5588 |
| ExoSAP-IT™ | Applied Biosystems™ | 78200 |
| 1X dsDNA High Sensitivity Kit, Qubit™ | Invitrogen™ | Q32851 |
| Native Barcoding Kit 96 V14 | Oxford Nanopore | NBD114.96 |
| R10.4.1 Flow Cell | Oxford Nanopore | FLO-MIN114 |
| Blunt/TA Ligase Master Mix | New England Biolabs | M0367 |
| NEBNext Ultra II End Repair/dA-Tailing Module | New England Biolabs | E7546 |
| NEBNext Quick Ligation Module | New England Biolabs | E6056 |
| Bovine Serum Albumin, UltraPure™ | Invitrogen™ | AM2616 |

### 3.0 Inhibitor Removal

For best practice, a process control negative and positive sample should be implemented from this point onwards. Norovirus GI and GII positive reference materials can be obtained from the UK Health Security Agency in the form of Virus LENTICULE® Discs. They will need dissolving in 975 µL PBS and then nucleic acid extraction prior to inputting into this process.

1. Pipette 25 µL of RNA extract into a 96 well PCR plate
2. Add 45 µL (1.8X) of Mag-Bind® Total Pure NGS beads
3. Pipette carefully 10x to mix
4. Leave to stand at room temperature for 5 min
5. If there is liquid on the side of the wells, cover the plate with a PCR adhesive seal and pulse-spin down in a plate centrifuge (with swing out rotor)
6. Place on magnetic rack for 3 min
7. Remove 68 µL of supernatant
8. Add 80 µL of 80% ethanol v/v (nuclease free water) for 30 sec (make EtOH fresh every time)
9. Remove 82 µL of supernatant being careful to avoid the pellet
10. Repeat clean once more (steps 8 and 9)
11. Set pipette to 20 µL and remove any residual ethanol
12. Leave to dry for 2 min
13. Remove from magnetic rack
14. Add 27 µL of molecular grade water and mix by pipetting 10x
15. Allow to stand for 5 min at room temperature
16. Place plate back on magnetic rack for 3 min
17. Recover 25 µL of the supernatant

### 4.0 Reverse Transcription

1. Add 2.5 µL of LunaScript® RT SuperMix to a 0.2 mL PCR reaction tube on ice on a cool block
2. Add 10 µL of cleaned nucleic acid extract and mix gently by pipetting 5x
3. Cover or cap the reaction tubes and briefly spin down
4. Incubate the combined reaction mixture with a heated lid (105°C) at:
    a. 25°C for 2 min
    b. 55°C for 45 min
    c. 95°C for 1 min
    d. Hold at 4°C.
5. Add 7.5 µL of molecular biology grade water to each sample
6. The first-strand cDNA can be stored in the fridge at 4 °C overnight or in the freezer at -20 °C if longer term storage is required.

### 5.0 Primer Preparation

1. Cartridge purified primers should be purchased to prevent loss of the degenerate oligonucleotides
2. Reconstitute the lyophilised primers in Table 2 to 100 µM using TE buffer pH8
3. Dilute primers to a working concentration of 10 µM in molecular biology grade water and aliquot into suitable volumes to prevent repeated freeze-thaw cycles

*Table 2. Norovirus GI and GII Primers*

| Name | Genogroup | F/R | Sequence (5’-3’) |
|---|---|---|---|
| $NV4478_m$ | GI | $F_1$ | AARYTVCCHATHAARGTTGGNATG |
| $NV4562_m$ | GI | $F_2$ | GATGCDGAYTAYACRGCHTGGG |
| $GISKR_m$ | GI | $R_{1,2}$ | CCIACCCAICCATTRTACA |
| NV4611 | GII | $F_1$ | CWGCAGCMCTDGAAATCATGG |
| NV4692 | GII | $F_2$ | GTGTGRTKGATGTGGGTGACTT |
| GIISKR | GII | $R_{1,2}$ | CCRCCNGCATRHCCRTTRTACAT |

Subscript represent **1**) first round primers and **2**) semi-nested primers

### 6.0 First-Round PCR

All PCR cycling conditions used a heated lid at 105°C and a ramping speed of 3°C/s. Prepare enough mastermix for all of the samples plus the process and PCR positive and negative controls. GI and GII PCRs are performed independently for each wastewater sample under analysis.

1. Prepare the mastermixes as indicated in Table 3 and Table 4 for the first round PCR
    a. Remove all mastermix components from the freezer and place the polymerase on ice
    b. Defrost all other components at room temperature
    c. Briefly vortex and spin down all components
    d. Add the components as outlined in Table 3 and 4 reserving the polymerase until the end
    e. When adding the polymerase, slowly aspirate and pre-wet pipette tip x3
    f. After dispensing the polymerase rinse the pipette tip 10x
    g. Vortex then briefly spin down the prepared mastermix
2. Distribute 20 µL of PCR mastermix 0.2 mL reaction tubes
3. Add 5 µL of cDNA for each sample
4. Add 5 µL molecular biology grade water or positive control cDNA for your PCR positive and negative controls
5. Seal or cap the reaction tubes then spin down ensuring no bubbles are present
6. Run the reactions in a thermocycler using the conditions in Table 5 and Table 6 for GI and GII

*Table 3. Forward and reverse primers for the GI and GII first-round PCRs*

| Genogroup | Forward Primer | Reverse Primer |
|---|---|---|
| GI | $NV4478_m$ | $GISKR_m$ |
| GII | NV4611 | GIISKR |

*Table 4. Mastermix Recipe for all PCRs*

| Reagents | Final concentration | Volume (µL) |
|---|---|---|
| $H_20$ (molecular biology grade) | - | 15.15 |
| PCR Buffer, without Mg (10X) | 1 | 2.5 |
| dNTP (10 mM) | 0.2 mM | 0.5 |
| $MgCl_2$ (50 mM) | 1.5 mM | 0.75 |
| Forward primer (10 µM) | 0.2 µM | 0.5 |
| Reverse primer (10 µM) | 0.2 µM | 0.5 |
| Platinum Taq polymerase (10 U/µL) | 1 unit/tube | 0.1 |

*Table 5. GI first-round PCR cycling conditions*

| Stage | Temperature (°C) | Time | Cycles |
|---|---|---|---|
| Initial denature | 95.0 | 60 s | 1 |
| Denature | 95.0 | 30 s | |
| Anneal | 47.4 | 30 s | 40 |
| Elongation | 72.0 | 30 s | |
| Final elongation | 72.0 | 7 min | 1 |
| Hold | 4.0 | ∞ | 1 |

*Table 6. GII first-round PCR cycling conditions*

| Stage | Temperature (°C) | Time | Cycles |
|---|---|---|---|
| Initial denature | 95.0 | 60 s | 1 |
| Denature | 95.0 | 30 s | |
| Anneal | 55.7 | 30 s | 40 |
| Elongation | 72.0 | 30 s | |
| Final elongation | 72.0 | 7 min | 1 |
| Hold | 4.0 | ∞ | 1 |

### 7.0 Semi-Nested PCR

Prepare enough mastermix for all of the samples and controls from the first-round PCR plus an additional PCR negative control for the semi-nested PCR.

1. Prepare the master mix as indicated in Table 4 and Table 7
    a. Remove all mastermix components from the freezer and place the polymerase on ice
    b. Defrost all other components at room temperature
    c. Briefly vortex and spin down all components
    d. Add the components as outlined in Table 3 and 4 reserving the polymerase until the end
    e. When adding the polymerase, slowly aspirate and pre-wet pipette tip x3
    f. After dispensing the polymerase rinse the pipette tip 10x
    g. Vortex then briefly spin down the prepared mastermix
2. Vortex then briefly spin down the prepared mastermix
3. Distribute 20 µL of PCR mastermix into a 96-well plate
4. Vortex and spin down plate from the first round PCR
5. Add 5 µL of first-round PCR product from to each of the sample wells
6. Add 5 µL of molecular biology grade water for your negative control
7. Seal or cap the plate then spin down ensuring no bubbles are present
8. Run using the cycles in Table 8 and Table 9 for GI and GII

*Table 7. GI and GII primer pairs for the semi-nested PCR*

| Genogroup | Forward Primer | Reverse Primer |
|---|---|---|
| GI | NV4562m | GISKRm |
| GII | NV4692 | GIISKR |

*Table 8. GI semi-nested PCR cycling conditions*

| Stage | Temperature (°C) | Time | Cycles |
|---|---|---|---|
| Initial denature | 95.0 | 60 s | 1 |
| Denature | 95.0 | 30 s | |
| Anneal | 57.2 | 30 s | 40 |
| Elongation | 72.0 | 30 s | |
| Final elongation | 72.0 | 7 min | 1 |
| Hold | 4.0 | ∞ | 1 |

*Table 9. GII semi-nested PCR cycling conditions*

| Stage | Temperature (°C) | Time | Cycles |
|---|---|---|---|
| Initial denature | 95.0 | 60 s | 1 |
| Denature | 95.0 | 30 s | |
| Anneal | 55.7 | 30 s | 40 |
| Elongation | 72.0 | 30 s | |
| Final elongation | 72.0 | 7 min | 1 |
| Hold | 4.0 | ∞ | 1 |

### 8.0 Analysis of PCR Controls

This step uses a TapeStation for PCR product analysis, but gel electrophoresis with 2% agarose tris-borate EDTA gel run at 125 V for 35 min with a 100 bp ladder (Promega, USA), can be used.

1. Remove ScreenTape from packaging, ensure there are no bubbles in any of the electrophoresis lanes, flicking ScreenTape to remove bubbles if necessary
2. Place ScreenTape into the TapeStation and allow to come to room temperature for 30 min
3. Load 10 µL of D5000 reagents into the 0.2 mL TapeStation reaction tubes
4. Add 1 µL of ladder into the tube representing position TapeStation position A1
5. Load 1 µL of your positive and negative PCR and process controls into the 0.2 mL TapeStation reaction tubes
6. Cap tubes, vortex and then spin down ensuring no bubbles are present
7. Place tubes into the machine and analyse the samples
8. All controls should give their expected results and appropriate actions taken if negative controls show contamination

### 9.0 PCR Product Clean-Up

This is performed for both the GI and GII semi-nested PCR products relating to each wastewater sample.

1. Remove ExoSAP-IT from the -20 °C freezer and allow to defrost on ice
2. Briefly vortex and spin down reaction tubes or plates from the semi-nested PCR
3. Briefly vortex and spin down ExoSap-IT
4. Aliquot 10 µL of semi-nested PCR product into a new reaction tube or plate
5. Add 4 µL of ExoSap-IT to each
6. Seal or cap reaction vessel
7. Incubate with a heated lid (105°C):
    a. 37°C for 15 min

   b. 80°C for 15 min
   c. Hold at 4°C

### 10.0 PCR Product Quantification

This is performed for both the GI and GII cleaned semi-nested PCR products.

1. Add 198 µL High-Sensitivity dsDNA Qubit™ reagents to Qubit tubes for each sample being quantified
2. Load 2 µL of cleaned, semi-nested PCR product onto the side of the tube ensuring residual nucleic acids aren't present on the exterior of the pipette tip
3. Add 190 µL High-Sensitivity dsDNA Qubit™ reagents to Qubit tubes for your standards
4. Load 10 µL of each standard
5. Gently tap tubes to allow PCR products to enter Qubit reagents
6. Cap tubes, vortex and briefly spin down
7. Check for presence of bubbles, flick tubes and spin down again if necessary
8. Incubate for 2 min and then take same measure with the Qubit™

### 11.0 GI and GII PCR Product Pooling

At the end of this process, for every sample being analysed, you will have a combined 200 fmol of GI and GII PCR products comprised of 85.3 and 114.7 fmol of GI and GII amplicons, respectively. The final volume will be 11.5 µL; ready for use directly in the end-prep for library preparation.

For samples which do not have sufficient PCR yield (200 fmol of combined GI and GII cleaned semi-nested PCR products) undiluted, cleaned GI and GII PCR products can be pooled and input into end-prep. Adjustment of the quantity of end-prepped DNA input into native barcoding can then be made by varying the sample volume from 0.75-3 µl, reducing the volume of water accordingly, to retain 10 fmol of input DNA.

1. Calculate the volume of water required to dilute 8 µL of the cleaned semi-nested GI and GII PCR products to 14.83 fmol/µL and 19.94 fmol/µL.
   a. For each sample, convert the GI and GII Qubit derived PCR product concentrations from ng/µL to fmol/µL by inputting into software such as the NEBioCalculator ensuring to enter the amplicon lengths of 1110 bp for GI and 971 bp for GII.
   b. Calculate the GI dilution factor by dividing the fmol/µL of the PCR product by 14.83
   c. Calculate the GII dilution factor by dividing the fmol/µL of the PCR product by 19.94
   d. Calculate the volumes of water required by subtracting 1 from the dilution factor and then multiplying by 8
2. Vortex and then briefly spin down the cleaned, semi-nested PCR products
3. Separately aliquot 8 µL of cleaned, semi-nested GI and GII PCR products into new reaction tubes
4. Add the volumes of water calculated in the previous steps to the GI and GII amplicons
5. Cap the reaction vessels
6. Briefly vortex and then spin down
7. In new reaction tubes, for every sample, combine 5.75 µL of both the diluted GI and GII PCR products for every sample undergoing analysis
8. Briefly vortex and then spin down

### 12.0 Native Barcoding and Sequencing

1. Perform end-prep and barcoding following the manufacturer’s instructions for ligation sequencing of amplicons in the Native Barcoding Kit 96 V14 by Oxford Nanopore Technologies
2. Load 45 fmol of library per flow cell, using an amplicon size of 1 kb for molarity calculation
3. Run sequencing at 260 bps using super-accurate basecalling until ≈60,000 reads per barcode have been generated

### 13.0 Set Up Software Environment

1. Install all the required software using mamba:

```
$ mamba create --name=amplion_analysis_norovirus duplex-tools=0.2.14 cutadapt=3.4 seqtk=1.3
minimap2=2.24 yacrd=1.0.0 kma=1.4.9 seqkit=2.3.0 samtools=1.13 bedtools=2.30.0 cd-hit=4.8.1
pyfastx=0.8.4

$ mamba create --name=NGSpeciesID medaka=1.2.2 bcftools=1.11 python=3.6.10 perl=5.32.0
openblas=0.3.3 spoa=4.0.7 racon=1.4.20 minimap2=2.17 tensorflow=2.4.1

$ mamba activate NGSpeciesID

$ pip install NGSpeciesID==0.1.3
```

2. Edit the NGSpeciesID script:

```
$ nano ~/mambaforge/envs/NGSpeciesID/bin/NGSpeciesID
```

3. Change the behaviour of the abundance_cutoff parameter so that it is based on the minimum coverage, rather than the relative abundance of a sequence within the sample:

```
-    abundance_cutoff = int( args.abundance_ratio * len(read_array))
+    abundance_cutoff = args.abundance_ratio
```

4. Edit NGSpeciesID's consensus module:

```
$ nano ~/mambaforge/envs/NGSpeciesID/lib/python3.6/site-packages/modules/consensus.py
```

5. Change the behaviour of NGSpeciesID so it is possible to output consensus sequences from spoa without polishing the consensus sequences using Medaka:

```
+    polishing = False
     if args.medaka:
         polishing_pattern = os.path.join(args.outfolder, "medaka_cl_id_*")
+        polishing = True
     elif args.racon:
         polishing_pattern = os.path.join(args.outfolder, "racon_cl_id_*")
+        polishing = True

-    for folder in glob.glob(polishing_pattern):
-        shutil.rmtree(folder)
+    if (polishing == True):
+        for folder in glob.glob(polishing_pattern):
+            shutil.rmtree(folder)

     spoa_pattern = os.path.join(args.outfolder, "consensus_reference_*")
     for file in glob.glob(spoa_pattern):
```

### 14.0 Split and Trim Reads

1. Create a folder to store results from the analysis:

```
$ mkdir Analysis_Results
```

```
$ cd Analysis_Results
```

2. Copy the fastq_pass or pass folder from the sequencing run to the Analysis_Results folder.
3. Store the name of the barcode and sample to be analysed as a variable:

```
$ barcodeID=barcode09

$ sampleID=Sample-4_PCR-Norm_PoolingType-1
```

4. Activate the correct software environment:

```
$ mamba activate amplion_analysis_norovirus
```

5. Use duplex_tools to split the reads:

```
$ mkdir Split_Reads

$ duplex_tools split_on_adapter --threads 12 --allow_multiple_splits \
  fastq_pass/${barcodeID} Split_Reads/${barcodeID} Native
```

6. Create a text file to store the sequence of different primers being used in the experiment:

```
$ nano primer_sequences.fasta
```

7. Copy the following into the file and then save it

```
>Long_GI_PCR1
AARYTVCCHATHAARGTTGGNATG...TGTAYAATGGNTGGGTNGG
>Long_GI_PCR2
GATGCDGAYTAYACRGCHTGGG...TGTAYAATGGNTGGGTNGG
>Long_GII_PCR1
CWGCAGCMCTDGAAATCATGG...ATGTAYAAYGGDYATGCNGGYGG
>Long_GII_PCR2
GTGTGRTKGATGTGGGTGACTT...ATGTAYAAYGGDYATGCNGGYGG
```

8. Use cutadapt to trim primer sequences from reads:

```
$ mkdir Trim_Reads

$ cat Split_Reads/${barcodeID}/*_split.fastq.gz >
Split_Reads/${barcodeID}/combined.fastq.gz

$ cutadapt -j12 --action=trim -n 1 -e 0.30 -O 12 --revcomp \
  -g file:primer_sequences.fasta --discard-untrimmed \
  --output Trim_Reads/trimmed-${sampleID}-{name}.fastq.gz \
  Split_Reads/${barcodeID}/combined.fastq.gz \
  > Trim_Reads/trimmed-${sampleID}.log 2>&1
```

9. Update the sampleID and barcodeID variables, and repeat steps described in this section for any other samples in the sequencing library.

### 15.0 Detect Chimeras

1. Select reads which include primers from the 2nd round of PCR and are longer than 800bp, then randomly sample 90,000 reads for subsequent filtering and chimera removal:

```
$ cat Trim_Reads/trimmed-${sampleID}-*_G*_PCR2.fastq.gz \
  > Trim_Reads/trimmed-${sampleID}-combined-PCR2.fastq.gz

$ seqtk seq -L 800 Trim_Reads/trimmed-${sampleID}-combined-PCR2.fastq.gz |\
  seqtk sample - 90000 > Trim_Reads/trimmed-${sampleID}-combined-PCR2-gt800bp.fastq
```

2. Use minimap2 to carry out an all-vs-all alignment of reads:

```
$ mkdir Detect_Chimeric_Reads
```

```
$ minimap2 -k19 -Xw19 -e0 -m100 -r100 -I 30M --cap-kalloc=8000m --cap-sw-mem=100m -t 12 \
  Trim_Reads/trimmed-${sampleID}-combined-PCR2-gt800bp.fastq \
  Trim_Reads/trimmed-${sampleID}-combined-PCR2-gt800bp.fastq \
  > Detect_Chimeric_Reads/trimmed-${sampleID}-combined-PCR2.overlap.paf
```

3. Use yacrd to identify chimeras and reads with poor support:

```
$ yacrd -t12 -c 10 -n 0.2 \
  -i Detect_Chimeric_Reads/trimmed-${sampleID}-combined-PCR2.overlap.paf \
  -o Detect_Chimeric_Reads/trimmed-${sampleID}-combined-PCR2.overlap-gt800bp.report.yacrd

$ awk '$1=="NotBad"' \
  Detect_Chimeric_Reads/trimmed-${sampleID}-combined-PCR2.overlap-gt800bp.report.yacrd |\
  cut -f2 \
  > Detect_Chimeric_Reads/trimmed-${sampleID}-combined-PCR2.overlap-gt800bp.report.lst

$ seqtk subseq Trim_Reads/trimmed-${sampleID}-combined-PCR2-gt800bp.fastq \
  Detect_Chimeric_Reads/trimmed-${sampleID}-combined-PCR2.overlap-gt800bp.report.lst \
  > Trim_Reads/trimmed-${sampleID}-combined-PCR2-gt800bp.filtered.fastq

$ rm  Detect_Chimeric_Reads/trimmed-${sampleID}-combined-PCR2.overlap.paf
```

4. Update the sampleID and barcodeID variables, and repeat the steps described in this section for any other samples in the sequencing library.

### 16.0 Creating Representative Sequences

1. Cluster reads to generate consensus sequences:

```
$ mkdir -p Consensus_Reads/${sampleID}/

$ mamba activate NGSpeciesID

$ NGSpeciesID --t 20 --q 15 --ont --consensus --max_seqs_for_consensus 100000 \
  --abundance_ratio 100 --m 1100 --s 200 \
  --fastq Trim_Reads/trimmed-${sampleID}-combined-PCR2-gt800bp.filtered.fastq \
  --outfolder Consensus_Reads/${sampleID}/ \
  > Consensus_Reads/${sampleID}/NGSpeciesID.log 2>&1 \
  || echo "No Assembly"
```

2. Align reads against consensus sequences, call variants and identify any poorly supported regions:

```
$ mamba activate amplicon_analysis_norovirus

$ cat Consensus_Reads/${sampleID}/consensus_reference_*.fasta \
> Consensus_Reads/${sampleID}_consensus_seqs.fasta

$ kma index -NI -i Consensus_Reads/${sampleID}_consensus_seqs.fasta \
  -o Consensus_Reads/${sampleID}_consensus_seqs

$ mkdir Align_vs_Consensus_Seqs

$ kma -i Trim_Reads/trimmed-${sampleID}-combined-PCR2.fastq.gz \
  -o Align_vs_Consensus_Seqs/${sampleID}_consensus_seqs_kma \
  -t_db Consensus_Reads/${sampleID}_consensus_seqs \
  -vcf 1 -ConClave 2 -bcNano -bc 0.7 -bcd 100 -t 20 -md 100 -1t1 \
  -ont -ml 900 -xl 1100 -ef -mrs 0.92 -mrc 0.90 \
  > Align_vs_Consensus_Seqs/${sampleID}_consensus_seqs_kma.log 2>&1
```

3. Update the sampleID and barcodeID variables, and repeat the last two steps for any other samples in the sequencing library.
4. Combine consensus sequences from every sample into a single file and rename them:

```
$ mkdir Cluster_Consensus_Sequences

$ cat Align_vs_Consensus_Seqs/*kma.fsa > Cluster_Consensus_Sequences/combined_seqs.fasta

$ seqtk rename Cluster_Consensus_Sequences/combined_seqs.fasta OTU_ \
  > Cluster_Consensus_Sequences/combined_seqs_renamed.fasta
```

5. Trim consensus sequences to remove any poorly supported regions identified by kma:

```
$ seqkit locate --bed -P -r -p '^[agct]+' -p '[agct]+$' \
  Cluster_Consensus_Sequences/combined_seqs_renamed.fasta \
  Cluster_Consensus_Sequences/combined_seqs_renamed.bed

$ samtools faidx Cluster_Consensus_Sequences/combined_seqs_renamed.fasta

$ bedtools sort -i Cluster_Consensus_Sequences/combined_seqs_renamed.bed \
  -g Cluster_Consensus_Sequences/combined_seqs_renamed.fasta.fai \
  > Cluster_Consensus_Sequences/combined_seqs_renamed_sorted.bed

$ bedtools complement -i Cluster_Consensus_Sequences/combined_seqs_renamed_sorted.bed \
  -g Cluster_Consensus_Sequences/combined_seqs_renamed.fasta.fai \
  > Cluster_Consensus_Sequences/combined_seqs_renamed_sorted_inverse.bed

$ bedtools getfasta -fi Cluster_Consensus_Sequences/combined_seqs_renamed.fasta \
  -bed Cluster_Consensus_Sequences/combined_seqs_renamed_sorted_inverse.bed \
  -fo Cluster_Consensus_Sequences/combined_seqs_renamed_trimmed.fasta
```

6. Cluster consensus sequences at 95 percent sequence identity:

```
$ cd-hit-est -i Cluster_Consensus_Sequences/combined_seqs_renamed_trimmed.fasta \
  -o Cluster_Consensus_Sequences/combined_seqs_renamed_trimmed_clustered.fasta \
  -G 0 -c 0.95 -n 10 -d 0 -M 100000 -T 80 -g 1 -aL 0.9
```

### 17.0 Calculating Coverage

1. Align reads against consensus sequences to calculate coverage:

```
$ mkdir Align_vs_Single_Set_Consensus_Reads

$ kma index -NI \
  -i Cluster_Consensus_Sequences/combined_seqs_renamed_trimmed_clustered.fasta \
  -o Cluster_Consensus_Sequences/combined_seqs_renamed_trimmed_clustered

$ kma -i Trim_Reads/trimmed-${sampleID}-combined-PCR2.fastq.gz \
  -o Align_vs_Single_Set_Consensus_Reads/${sampleID}_consensus_seqs_kma \
  -t_db Cluster_Consensus_Sequences/combined_seqs_renamed_trimmed_clustered \
  -nc -vcf 2 -ConClave 2 -bcNano -bc 0.7 -bcd 100 -t 20 -md 100 -1t1 \
  -ont -ml 900 -xl 1100 -ef -mrs 0.92 -mrc 0.90 \
  > Align_vs_Single_Set_Consensus_Reads/${sampleID}_consensus_seqs_kma.log 2>&1
```

2. Use the following files for downstream statistical analysis:

The coverage of each representative sequence in each sample:

```
Align_vs_Single_Set_Consensus_Reads/*_consensus_seqs_kma.mapstat
```

The sequence of each representative sequence:

```
Cluster_Consensus_Sequences/combined_seqs_renamed_trimmed_clustered.fasta
```

### 18.0 Collating Coverage, Sequencing and Typing Results

1. Use the CDC's calicivirus typing tool to type each representative sequence:
2. https://calicivirustypingtool.cdc.gov/
3. Upload the entire fasta file
4. Click on 'type it'
5. Once the input sequences have been typed click on 'Download EXCEL'
6. Install R (≥ 4.1.2), RStudio (≥ 1.4.1717) and the openxlsx package (≥ 4.2.5.2)
7. Setup a new project directory using RStudio
8. Download the coverage files to a sub-folder within the project directory called 'Coverage Results'

9. Copy the Excel file downloaded from the CDC's calicivirus typing tool website to the project directory
10. Create a new R script in your RStudio project
11. Copy the following code into the R script:

```
# Load library for opening xlsx files
library(openxlsx)

# List coverage files
coverage_results =
  list.files(path = 'Coverage Results', pattern = "\\_consensus_seqs_kma.mapstat",
             recursive = TRUE, include.dirs = FALSE)

# List samples
samples = gsub(x = coverage_results,
               pattern = "^(.+)_consensus_seqs_kma.mapstat", replacement = "\\1")

# Collate coverage results into a single file
combined_coverage_results = NULL

for (sampleID in 1:length(samples)) {
  print(sampleID)
  coverage_filename =
    paste0("Coverage Results/", samples[sampleID], "_consensus_seqs_kma.mapstat")

  coverage_file = readLines(con = coverage_filename)

  coverage_file[[7]] = gsub(x = coverage_file[[7]], pattern = "# ", replacement = "")

  noRecords = sum(!grepl(pattern = "##", x = coverage_file,fixed = TRUE)) - 1

  if (noRecords > 0) {
    coverage = read.table(file = textConnection(coverage_file),
                          header = TRUE, stringsAsFactors = FALSE, sep = "\t", skip = 6)

    coverage$sample = samples[sampleID]

    combined_coverage_results = rbind(combined_coverage_results, coverage)
  }

}

# Import file linking reference sequences to genotype
taxa_results_field_names =
  c('query','name','sequence','length','Genus','Genotype-B-region-score',
    'Genotype-B-region-plot','Genotype','Genotype-C-region-plot',
    'Genotype-C-region-score','Genotype-plot')

combined_taxa_results =
  read.xlsx(xlsxFile = "calicivirus_typing_output.xlsx",
            sheet = "sheet1", startRow = 2)

colnames(combined_taxa_results) = taxa_results_field_names

# Merge coverage results with genotype
combined_coverage_results2 =
  merge(x = combined_coverage_results,
        y = combined_taxa_results[,c('name','Genotype')],
        by.x = c('refSequence'), by.y = c('name'), all.x = TRUE)

combined_coverage_results2[
  is.na(combined_coverage_results2$Genotype),
  c('Genotype')
  ] = "Unknown"

# Summarise by genotype
coverage_per_genotype =
  aggregate(formula = readCount ~ Genotype + refSequence + sample,
            data = combined_coverage_results2,
            FUN = sum)

coverage_per_genotype_wide =
  reshape(data = coverage_per_genotype,
          v.names = "readCount",
```

```
          idvar = c("Genotype","refSequence"),
          timevar = "sample", direction = "wide")

coverage_per_genotype_wide[is.na(coverage_per_genotype_wide)] = 0

colnames(coverage_per_genotype_wide) =
  gsub(x = colnames(coverage_per_genotype_wide),
       pattern = "^readCount\\.(.+)$", replacement = "\\1")

# Save results as a csv file
write.csv(x = coverage_per_genotype_wide,
          file = "coverage_per_genotype_wide.csv", row.names = FALSE)
```

12. Run the R script, check for errors and that the output looks correct

### 19.0 Read Depth Filtering

1. Using the coverage_per_genotype_wide.csv file generated above, for GI and GII independently, sum the norovirus-aligned reads for each sample
2. Find the median of the total aligned reads per sample across the data set
3. Filter data on a sample-to-sample basis removing reads associated with consensus sequences that have fewer than 0.1% of the median total aligned reads per sample

### 20.0 Reference Database Removal of PCR Chimeras

PCR chimeras should be present at relatively low abundances. It is worth taking note of the No. of reads aligned with any putative chimeras to avoid the removal of novel recombinants without further investigation.

1. Collate a list of all known types of norovirus GI and GII from the Centre for Disease Control's Human Calicivirus Typing Tool – this should be performed prior the analysis of any new data sets to prevent missing recently observed types
2. Remove all coverage (coverage_per_genotype_wide.csv) and sequence data (combined_seqs_renamed_trimmed_clustered.fasta) relating to consensus sequences that are types not previously reported by the CDC.
    a. This can be done manually for small data sets or using R and tidyverse 2.0.0

### 21.0 De Novo Removal of PCR Chimeras

1. For each wastewater sample under analysis, annotate the consensus sequences (combined_seqs_renamed_trimmed_clustered.fasta) with the read depths (coverage_per_genotype_wide.csv)
    a. This can be done manually or with R using seqinr 4.2.3 and tidyverse 2.0.0
2. Perform chimera filtering with USEARCH v11 using the following parameters:
    a. -uchime3_denovo -chimeras chimeras.fa
3. Remove all coverage (coverage_per_genotype_wide.csv) and sequence data (combined_seqs_renamed_trimmed_clustered.fasta) relating to consensus sequences identified as chimeras in chimeras.fa

### 22.0 Manual Screening for PCR Chimeras

1. For each sample undergoing analysis, identify types that may be generated from two other (parent) types in the sample using the data in coverage_per_genotype_wide.csv.
    a. For example, GI.6[P13] may be a chimera of GI.6[P11] and GI.3[P13] if the latter two have a higher number of reads associated with them

2. Assess the putative chimeras and parent sequences using a multiple sequence alignment tool such as NCBI Multiple Sequence Alignment Viewer v1.25.0 with the putative chimera sequence anchored
3. Check the parent sequences for breakpoints within the terminal or proximal regions of *RdRp* and *VP1* and child-parent sequence similarities ≥95%
4. Remove all coverage (coverage_per_genotype_wide.csv) and sequence data (combined_seqs_renamed_trimmed_clustered.fasta) relating to consensus sequences identified as putative chimeras which match these criteria
5. Your data is now ready for downstream analysis. It should be noted, however, that at present this method is only suitable for qualitative analysis and read-depth/coverage should not be used as a quantitative measurement of the different types detected in a sample
